# Ontario Neurodegenerative Disease Research Initiative (ONDRI): Structural MRI methods & outcome measures

**DOI:** 10.1101/2019.12.13.875823

**Authors:** Joel Ramirez, Melissa F. Holmes, Christopher J.M. Scott, Miracle Ozzoude, Sabrina Adamo, Gregory M. Szilagyi, Maged Goubran, Fuqiang Gao, Stephen R. Arnott, Jane M. Lawrence-Dewar, Derek Beaton, Stephen C. Strother, Douglas P. Munoz, Mario Masellis, Richard H. Swartz, Robert Bartha, Sean Symons, Sandra E. Black, the ONDRI Investigators

## Abstract

The Ontario Neurodegenerative Research Initiative (ONDRI) is a 3 year multi-site prospective cohort study that has acquired comprehensive multiple assessment platform data, including 3T structural MRI, from neurodegenerative patients with Alzheimer’s disease, mild cognitive impairment, Parkinson’s disease, amyotrophic lateral sclerosis, frontotemporal dementia, and cerebrovascular disease patients. This heterogeneous cross-section of patients with complex neurodegenerative and neurovascular pathologies pose significant challenges for standard neuroimaging tools. To effectively quantify regional measures of normal and pathological brain tissue volumes, the ONDRI neuroimaging platform implemented a semi-automated MRI processing pipeline that was able to address many of the challenges resulting from this heterogeneity. This paper describes the comprehensive neuroimaging pipeline methods used to generate regional brain tissue volumes & neurovascular markers.

## 1. INTRODUCTION

The Ontario Neurodegenerative Research Initiative (ONDRI) is a multi-site prospective cohort study following patients with neurodegenerative diseases including Alzheimer’s disease (AD), mild cognitive impairment (MCI), Parkinson’s disease (PD), amyotrophic lateral sclerosis (ALS), frontotemporal dementia (FTD), and cerebrovascular disease (CVD) (Farhan et al., 2017). Over the course of 3 years, multiple assessment platforms acquired comprehensive data from the 520 patients including, clinical and demographic assessments, neuroimaging, neuropsychology, genomics, eye tracking and pupillometry, thickness of retinal layers, gait performance, and neuropathology (Dilliott et al., 2019; Farhan et al., 2017; Wong et al., 2019). The multi-modal data collected from ONDRI will be used to explore earlier detection, guide development of novel therapy, and improve patient care (Sunderland et al., 2019).

This paper describes the methods implemented to extract normal and pathological brain tissue volumetric information from the structural Magnetic Resonance Imaging (MRI) provided by the ONDRI neuroimaging platform. It includes a comprehensive methodological overview of the structural neuroimaging pipeline components, with numerous figures to provide a visual description of how the measures were obtained from the MRI, some recommendations for reporting and data analysis, and a brief section providing some basic descriptive statistics to illustrate the whole brain volumetrics that can be obtained from the ONDRI patient cohorts.

Structural MRI processing for volumetrics was performed by the neuroimaging group (BrainLab.ca) in the L.C. Campbell Cognitive Neurology Research Unit, within the Hurvitz Brain Sciences Research Program, at the Sunnybrook Research Institute, in Toronto, Canada. The image processing pipeline (**Fig.1**) has been optimized for an aging population, with a particular emphasis on accounting for chronic stroke and post stroke cortical and subcortical lesions, numerous imaging markers of cerebral small vessel disease, as well as, the focal and global brain atrophy observed in neurodegenerative patient populations such as AD and FTD. For more information on the ONDRI project, please visit: http://ondri.ca/. Anonymized structural imaging data will be made publicly available in the future through an application process.

**Fig. 1.**
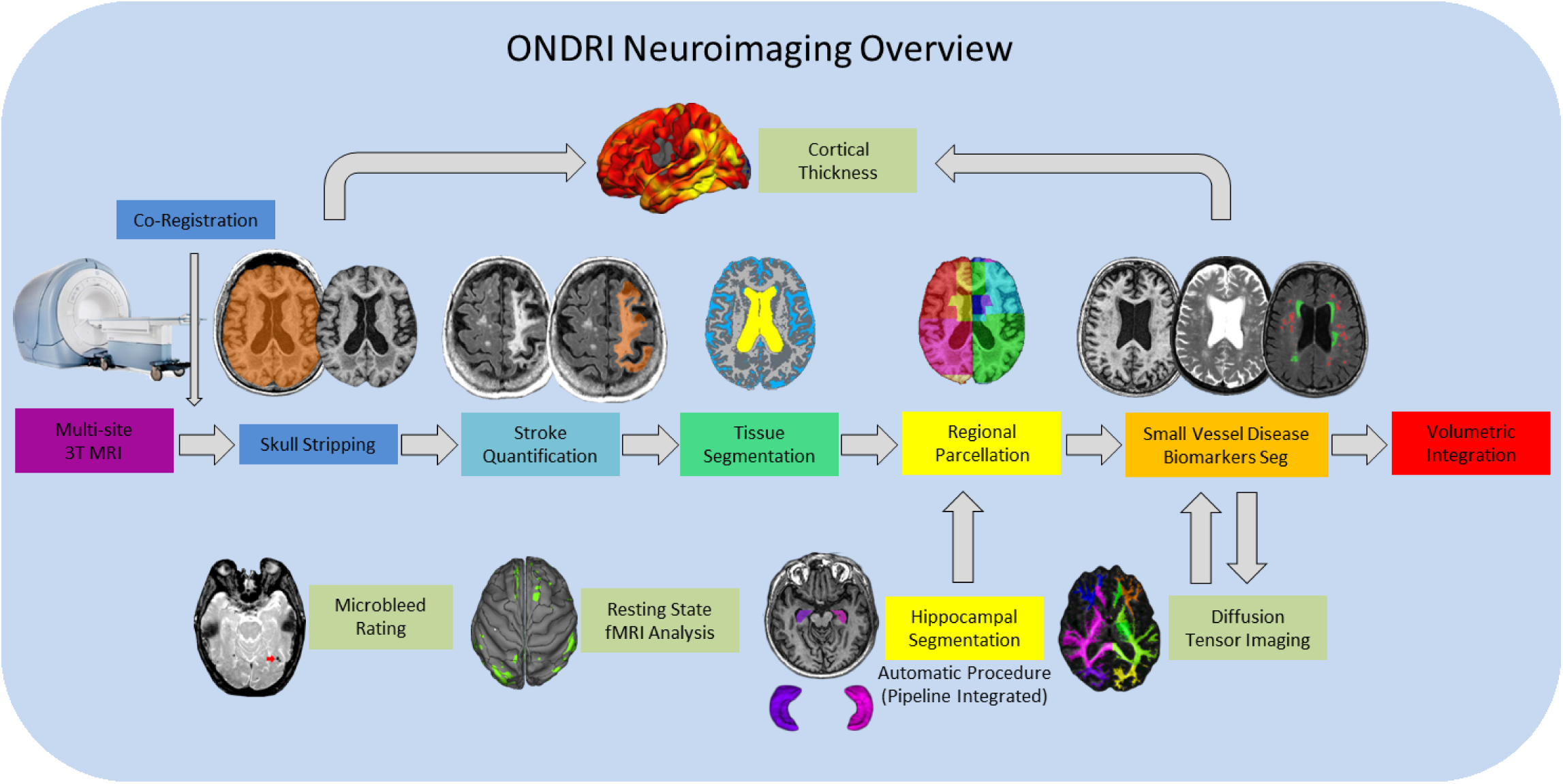
ONDRI MRI processing pipeline overview. General workflow moves from left to right for final volumetric output resulting in a comprehensive spreadsheet in the form of a .csv file. Hippocampal volumes are segmented using the SBHV method (Nestor et al., 2013) which was fully integrated into the pipeline and are included in the final volumetric spreadsheet. Microbleed Rating, Resting State fMRI Analysis, and Diffusion Tensor Imaging (DTI) analyses are processed separately, however, the DTI pipeline (Haddad et al., 2020) and Cortical Thickness pipelines are dependent on some components of the primary pipeline, thus, results from these processes are also provided in separate spreadsheets.

## 2. METHODS

### 2.1 Study Participants

Ethics approval was obtained from all participating institutions. Participants were recruited at 14 health centres across 6 cities in Ontario, Canada: Hamilton General Hospital and McMaster Medical Centre in Hamilton; Hotel Dieu Hospital and Providence Care Hospital in Kingston; London Health Science Centre and Parkwood Institute in London; Elizabeth Bruyère Hospital and The Ottawa Hospital in Ottawa; Thunder Bay Regional Health Sciences Centre in Thunder Bay; and Baycrest Health Sciences, Centre for Addiction and Mental Health, St. Michael’s Hospital, Sunnybrook Health Sciences Centre, and Toronto Western Hospital in Toronto.

Full study participant details are previously described (Farhan et al., 2017). Briefly, AD/MCI patients met National Institute on Aging Alzheimer’s Association criteria for probable or possible AD, or MCI (Albert et al., 2011; McKhann et al., 2011); PD patients met criteria for idiopathic PD defined by the United Kingdom’s Parkinson’s Disease Society Brain Bank clinical diagnostic criteria (Gibb and Lees, 1988); ALS patients met El Escorial World Federation of Neurology diagnostic criteria for possible, probable, or definite familial or sporadic ALS (Brooks, 1994); FTD patients included possible or probable behavioural variants of frontotemporal degeneration (Rascovsky et al., 2011), agrammatic/non-fluent and semantic variants of primary progressive aphasia (Gorno-Tempini et al., 2011), and possible or probable progressive supranuclear palsy (Höglinger et al., 2017); CVD patients experienced a mild to moderate ischemic stroke event, verified on neuroimaging, three or more months prior to enrollment in compliance with the National Institute of Neurological Disorders and Stroke-Canadian Stroke Network vascular cognitive impairment harmonization standards (Hachinski et al., 2006).

For illustrative purposes of the neuroimaging pipeline outputs, baseline MRI data are included for the following ONDRI patient cohorts: 126 AD/MCI, 140 PD, 40 ALS, 53 FTD, and 161 CVD.

### 2.2 MRI acquisition

Neuroimaging was acquired at the following sites using each respective 3T MRI system: a General Electric (GE, Milwaukee, WI) Discovery 750 was used at Sunnybrook Health Sciences Centre, McMaster University/Hamilton General Hospital, and the Centre for Addiction and Mental Health; a GE Signa HDxt at Toronto Western Hospital; a Philips Medical Systems (Philips, Best, Netherlands) Achieva system at Thunder Bay Regional Health Sciences Centre; a Siemens Health Care (Siemens, Erlangen, Germany) Prisma at Sunnybrook Health Sciences Centre and London Health Sciences Centre/Parkwood Hospital; a Siemens TrioTim at Ottawa Hospital/Élisabeth Bruyère Hospital, Hotel Dieu Hospital/Providence Care Hospital and Baycrest Health Sciences; and a Siemens Skyra at St. Michael’s Hospital.

Harmonized with the Canadian Dementia Imaging Protocol (Duchesne et al., 2019), the National Institute of Neurological Disorders and Stroke–Canadian Stroke Network Vascular Cognitive Impairment Harmonization Standards (Hachinski et al., 2006), full MRI acquisition protocol details for each imaging site are provided on **Supplementary Table 1**. In brief, the following structural MRI sequences were obtained for each study participant: 3D T1-weighted (T1), T2-weighted fluid attenuated inversion recovery (FLAIR), interleaved T2-weighted and proton density (T2/PD), and T2*gradient recalled echo (GRE). It should be noted that additional imaging protocol included a 30/32 direction diffusion tensor imaging (DTI). resting state functional MRI, and arterial spin labelling (acquired only at one site), but are beyond the scope of this paper and will be presented elsewhere (Hadadd et al., 2020). Prior to image processing for volumetric quantification, MRI were fully evaluated by a neuroradiologist (S.S.) for incidental findings and for imaging quality by a medical biophysics scientist (R.B.).

**Table 1.** SABRE-LE neuroimaging pipeline brain tissue and lesion codes (top), and detailed SABRE brain region codes (bottom). Note that each regional code will be preceded by an ‘L’ or ‘R’ indicating the left or right hemisphere.

| Imaging Description | Code | Section Number |
| --- | --- | --- |
| Supratentorial Total Intracranial Volume | ST_TIV | 2.5 |
| Normal Appearing Gray Matter | NAGM | 2.6 |
| Normal Appearing White Matter | NAWM | 2.6 |
| Sulcal Cerebrospinal Fluid | sCSF | 2.6 |
| Ventricular Cerebrospinal Fluid | vCSF | 2.6 |
| Periventricular White Matter Hyperintensities | pWMH | 2.7 |
| Deep White Matter Hyperintensities | dWMH | 2.7 |
| Periventricular Lacunes | pLACN | 2.8 |
| Deep Lacunes | dLACN | 2.8 |
| Enlarged Perivascular Spaces | PVS | 2.9 |
| Chronic Stroke Lesions | Stroke | 2.11 |
| SABRE Brain Region Name | Code | Lobe |
| Superior Frontal | SF | Frontal |
| Middle Frontal | MF | Frontal |
| Inferior Frontal | IF | Frontal |
| Medial Inferior Frontal | MIF | Frontal |
| Medial Superior Frontal | MSF | Frontal |
| Medial Middle Frontal | MMF | Frontal |
| Superior Parietal | SP | Parietal |
| Inferior Parietal | IP | Parietal |
| Occipital | O | Occipital |
| Anterior Temporal | AT | Temporal |
| Posterior Temporal | PT | Temporal |
| Anterior Basal Ganglia/Thalamus | ABGT | Basal Ganglia/Thalamus |
| Posterior Basal Ganglia/Thalamus | PBGT | Basal Ganglia/Thalamus |
| Hippocampus | HP | Medial Temporal |

### 2.3 Structural Image Processing Methods: Overview

The structural neuroimaging pipeline used in ONDRI is a component based algorithm commonly referred to as SABRE-Lesion Explorer (SABRE-LE) (Dade et al., 2004; Gibson et al., 2010; Ramirez et al., 2014, 2011). This is a semi-automated personalized approach to imaging-based quantification, as it can provide a comprehensive volumetric profile at the individual patient level. While it may take longer to process each individual relative to fully automatic methods, this careful patient-focused approach is more robust to the large variability in stroke and neurodegenerative patient population. This method has been previously validated (Kovacevic et al., 2002; Ramirez et al., 2013, 2011) and implemented in other Canadian studies (Dey et al., 2019; Ramirez et al., 2018; Sam et al., 2016; Swardfager et al., 2018). For additional information on the pipeline used to generate volumetrics outlined in this document, please visit: http://imaging.brainlab.ca/ The following sections describe the SABRE-LE comprehensive pipeline method and the volumetric data that is extracted in greater detail. Data visualization was performed using RStudio version 1.2.1335 (RStudio, Inc., Boston, MA) and ITKSnap (Yushkevich et al., 2006).

### 2.4. Brain Regions of Interest: SABRE

The neuroimaging pipeline integrates a brain region parcellation process called Semi-Automatic Brain Region Extraction (SABRE) (Dade et al., 2004). This method separates the brain into 28 regions of interest (ROIs: 13 per hemisphere + 2 hippocampi) derived from anatomical landmarks manually identified per hemisphere on each individual patient (**Fig. 2** and **Table 1**). Each imaging analyst was required to achieve an intraclass correlation coefficient (ICC) > 0.90 in order to work on ONDRI patient imaging analysis. The automatic SunnyBrook Hippocampal Volumetry (SBHV) tool (Nestor et al., 2013) was subsequently integrated into the SABRE pipeline (**Fig.3**), resulting in a total of 28 ROIs (left + right hippocampus) (see following section). The SABRE brain maps are personalized maps that are unique to each individual patient and was developed from the Talairach grid system (Talairach and Tournoux, 1988). Relative to many brain mapping methods that implement nonlinear (i.e. ‘warping’) techniques to register an individual patient’s MRI to a standardized template, such as the Montreal Neurological Institute brain (MNI152) (Mazziotta et al., 1995), the SABRE approach is essentially reversed, by mapping a brain template onto the individual patient’s MRI. This method accounts for natural individual differences in anatomy but more importantly, it is a method that can compensate for significant focal and global brain atrophy that is found in stroke, dementia and neurodegenerative patients.

**Fig. 2.**
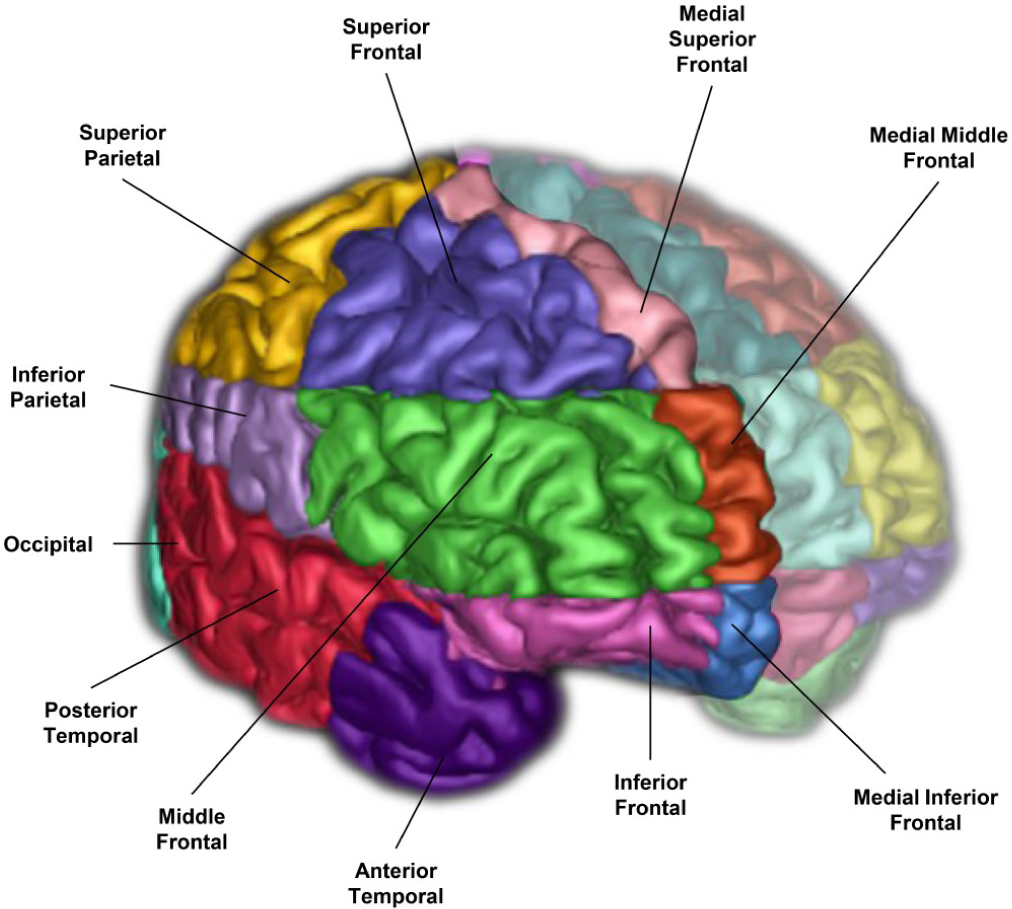
A 3-D surface volume rendering of T1-weighted MRI showing right hemisphere SABRE regions in different colours. Left hemi-sphere regions were made translucent for illustrative purposes, however, SABRE regions are separately parcellated for each hemisphere and delineated using individualized anatomical landmarks for both left and right sides.

**Fig. 3.**
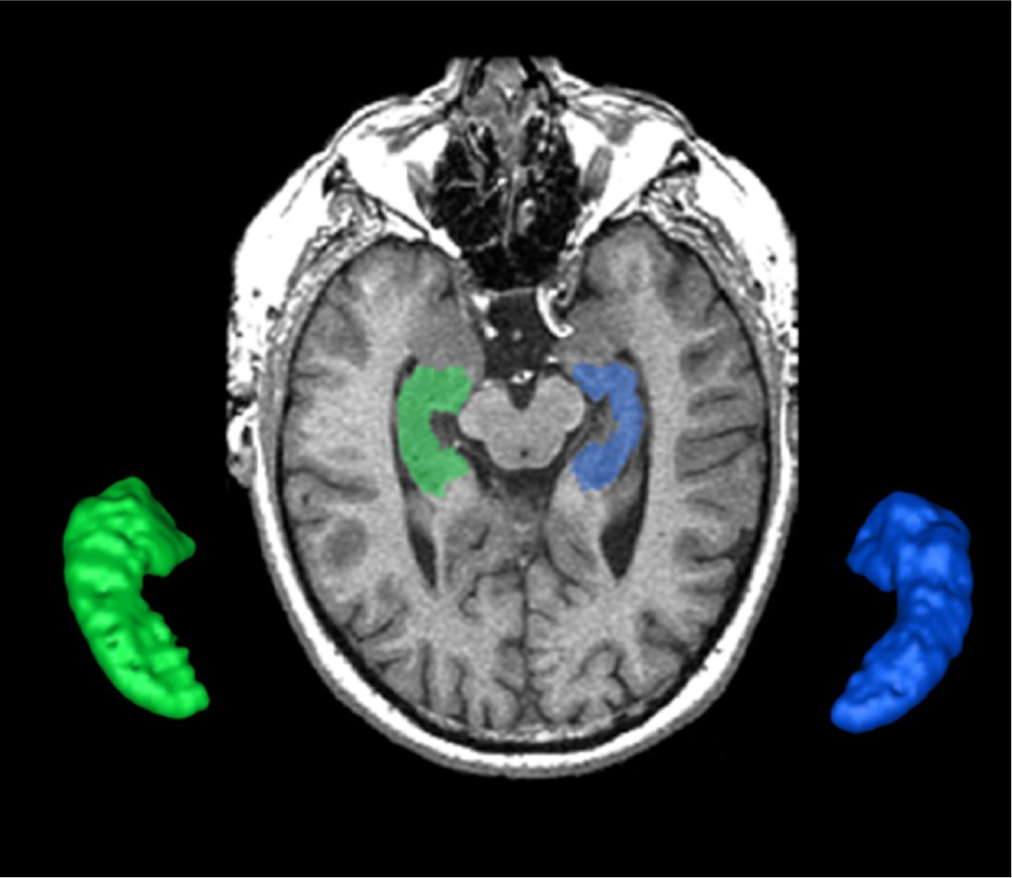
The SunnyBrook Hippocampal Volumetric (SBHV) segmentation showing left (BLUE) and right (GREEN) hippocampi over-layed on an axial T1 MRI and extracted as 3-D surface volume renderings. Note images are in radiological convention.

#### 2.4.1. Hippocampus

The hippocampus is an important part of the limbic system that has been studied extensively in dementia, given its significant role in memory functions (Frisoni and Jack, 2011; Moodley and Chan, 2014). The ONDRI pipeline incorporates the multi-atlas based Sunnybrook Hippocampal Volumetric (SBHV) segmentation tool (**Fig.3**) that was developed and validated using the Sunnybrook Dementia Study and the Alzheimer’s Disease Neuroimaging Initiative (ADNI1) (Nestor et al., 2013).

For ONDRI, the SBHV segmentation has been fully integrated into the SABRE-LE pipeline, and includes left and right hippocampal sub-classifications for parenchyma, hypointensities, and stroke volumes (when present). Currently, there is some controversy over the pathophysiological origin and relevance of small cavities commonly observed in the hippocampus (Bastos-Leite et al., 2006; Maller et al., 2011; Van Veluw et al., 2013; Yao et al., 2014), which are particularly relevant in the ONDRI CVD patients. Additionally, large cortico-subcortical strokes can extend from the cortex into the hippocampus. Given these vascular issues potentially affecting the overall hippocampal volume, ONDRI provides sub-classifications for parenchyma, hypointensities, and stroke volumes based on the neuroimaging characteristics (i.e. intensity) using the voxel segmentation classifications and takes a neutral stance on the pathophysiological origin of small cavities observed in this region.

### 2.5. Total Intracranial Volume

The supratentorial total intracranial volume (ST-TIV) is a measure of all brain matter that is located below the dura mater. It is referred to as *supratentorial* because the SABRE-LE method removes all tissue below the tentorium, including the cerebellum and portions of the brain stem (Dade et al., 2004; Ramirez et al., 2014).

In addition to sex-related differences, there are also normal variations in head size. In order to account for these differences, most neuroimaging studies implement some form of head-size correction. This is also particularly important when assessing brain atrophy in cross-sectional studies, as a true measure of the total intracranial capacity will provide an indication of where ‘there used to be brain and now there is cerebrospinal fluid (CSF)’. The presence of focal atrophy due to stroke and neurodegenerative processes tends to result in over and under erosion errors with many fully automated T1-based skull stripping techniques, due to the similarity in intensity between background and sulcal CSF. The SABRE-LE method accounts for the presence of focal atrophy since it includes a measure of everything below the dura mater, including sub-arachnoid, thus, providing a more accurate measure of head-size in neurodegenerative patient populations (**Fig.4**).

**Fig. 4.**
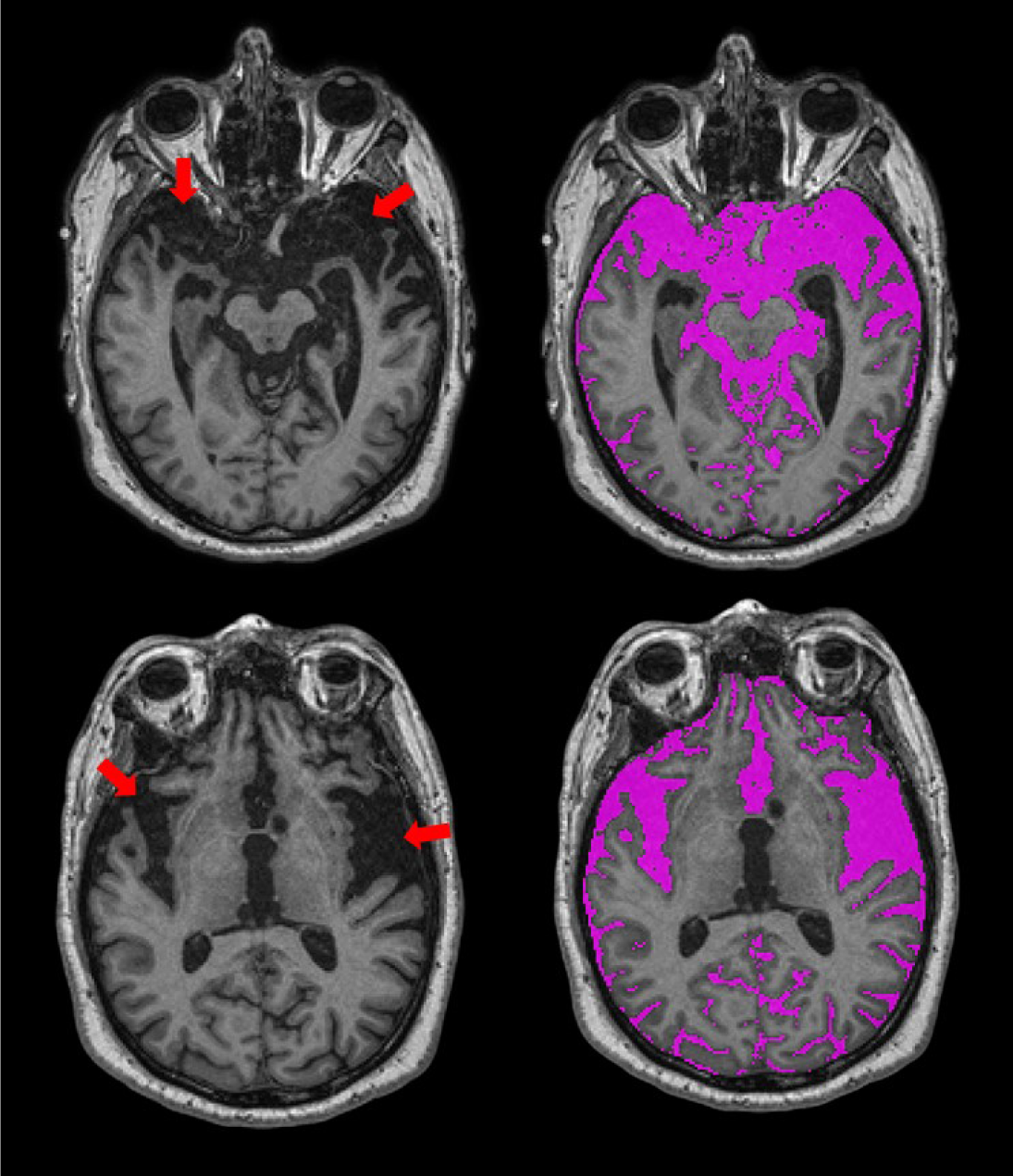
Axial views of T1-weighted MRI from an ONDRI FTD patient. Red arrows point to regions with significant focal brain atrophy. The SABRE-LE processing pipeline accounts for this focal atrophy since it includes a measure of everything below the dura mater, including sub-arachnoid and sulcal cerebrospinal fluid (CSF), shown in purple.

It is important to note that there are numerous acceptable head-size correction methods reported in the literature (Vågberg et al., 2017). A simple method involves dividing each volume of interest by the total head size to obtain a proportional volume (Rudick et al., 1999). ONDRI provides raw volumes and head size volumes (i.e., ST-TIV) for each individual patient.

### 2.6 Brain Tissue Segmentation

A robust T1 intensity-based brain tissue segmentation, optimized for aging and dementia, is performed after skull stripping and removal of non-brain tissue (Kovacevic et al., 2002). This automatic segmentation method deals with scanner inhomogeneities by fitting localized histograms to Gaussians to allocate voxels into gray matter (GM), white matter (WM) and cerebrospinal fluid (CSF) tissue classes. After manual ventricular CSF (vCSF) relabelling, there are 4 brain tissue types that are segmented for volumetrics using SABRE-LE (**Table 2**):

**Table 2.** Data is shown as mean (SD) unless otherwise specified. Raw values are presented for transparency purposes. *Data is shown as median (interquartile range); ^a^ available in 1/40 participants; ^b^ available in 6/52 participants; ^c^ available in 4/140 participants; ^d^ available in 88/155 participants. Abbreviations: AD/MCI = Alzheimer’s Disease & Mild Cognitive Impairment; ALS = Amyotrophic Lateral Sclerosis; FTD = Frontotemporal Dementia; PD = Parkinson’s Disease; CVD = Cerebrovascular Disease; ST-TIV = supratentorial total intracranial volume; NAWM = normal appearing white matter; NAGM = normal appearing gray matter; sCSF = sulcal cerebrospinal fluid; vCSF = ventricular cerebrospinal fluid; pWMH = periventricular white matter hyperintensities; dWMH = deep white matter hyperintensities; LACN = lacunes; PVS = perivascular spaces.

| Demographics | AD/MCI | ALS | FTD | PD | CVD |
| --- | --- | --- | --- | --- | --- |
| Number of participants | 126 | 40 | 52 | 140 | 155 |
| Age, years | 71.0 (8.2) | 62.0 (8.7) | 67.8 (7.1) | 67.9 (6.3) | 69.3 (7.4) |
| Sex, n (%) female | 57 (45.2) | 16 (40.0) | 19 (36.5) | 31 (22.1) | 48 (31.0) |
| ST-TIV, cc | 1235.6 (144.6) | 1203.6 (162.8) | 1245.8 (129.6) | 1316.6 (127.0) | 1224.5 (133.2) |
| NAWM, cc | 395.4 (344.5) | 425.0 (78.8) | 295.1 (59.4) | 446.1 (61.2) | 387.4 (54.4) |
| NAGM, cc | 533.3 (51.4) | 556.2 (65.7) | 252.5 (56.0) | 574.7 (47.1) | 535.7 (52.3) |
| sCSF, cc | 256.3 (62.1) | 195.9 (52.9) | 277.0 (57.8) | 252.3 (53.3) | 242.6 (59.3) |
| vCSF, cc | 45.7 (28.4) | 23.8 (11.1) | 43.7 (16.6) | 38.2 (19.4) | 41.3 (23.0) |
| pWMH*, mm<sup>3</sup> | 2564.5 (2811.2) | 1040.0 (1252.5) | 2736.0 (1623.8) | 2563.5 (2708.0) | 4054.0 (7468.0) |
| dWMH*, mm<sup>3</sup> | 289.5 (424.7) | 208.0 (386.5) | 138.5 (379.3) | 259.5 (225.7) | 555.0 (584.0) |
| LACN*, mm<sup>3</sup> | 15.5 (66.0) | 14.5 (12.2) | 9.5 (55.5) | 17.5 (70.0) | 92.0 (291.0) |
| PVS*, mm<sup>3</sup> | 45.5 (35.5) | 17.5 (9.5) | 32.5 (36.3) | 34.0 (30.0) | 44.0 (33.0) |
| Stroke*, mm<sup>3</sup> | -- | 90.0<sup>a</sup> | 393.0 (294.0)<sup>b</sup> | 531.5 (1269.0)<sup>c</sup> | 4644.5 (12963.0)<sup>d</sup> |

Normal appearing gray matter (NAGM)

Normal appearing white matter (NAWM)

Sulcal cerebrospinal Fluid (sCSF)

Ventricular CSF (vCSF)

The T1-based tissue segmentation is further corrected for misclassified volumes using a PD-T2/FLAIR-based lesion segmentation algorithm to account for the voxels appearing as GM or CSF on T1 (Levy-Cooperman et al., 2008) due to WM changes from stroke and cerebral small vessel disease. For this reason, the GM and WM volumes are denoted as ‘normal appearing’ (NAGM, NAWM) to signify that these volumes have been re-labelled as normal appearing after having been corrected with an additional multi-modal MRI segmentation approach (**Fig.5**). Additional brain tissue volumes for stroke lesions and cerebral small vessel disease markers are discussed in the following sections.

**Fig. 5.**
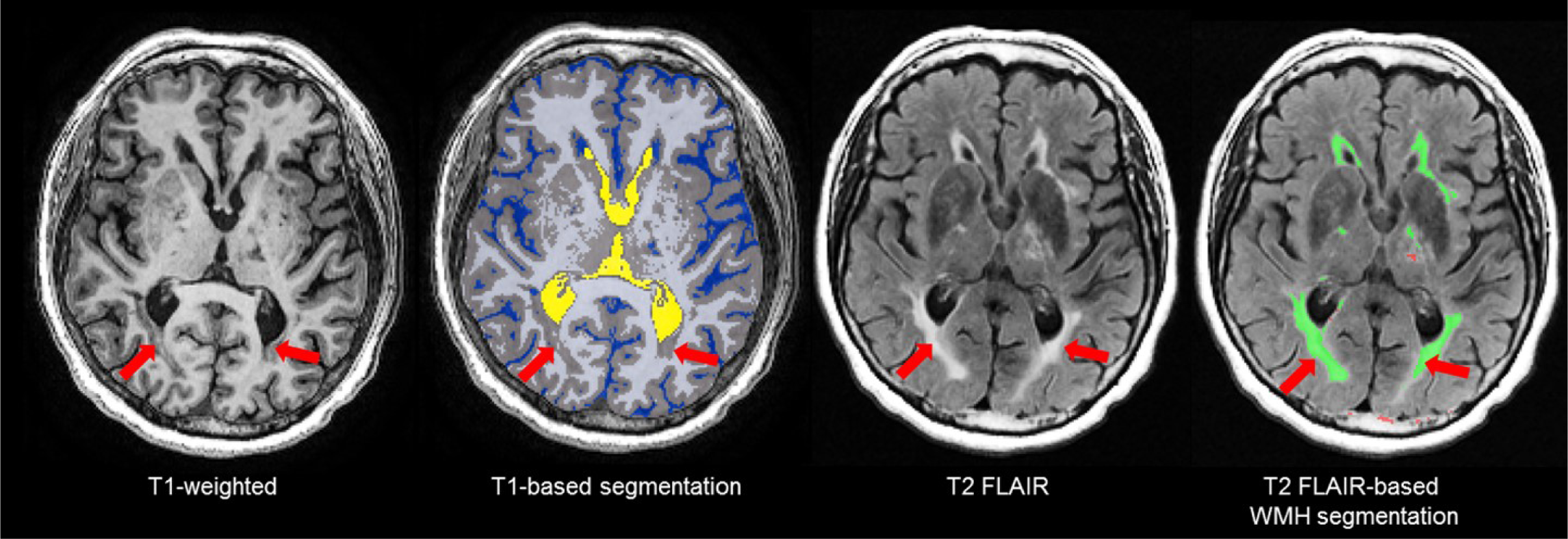
Due to relative intensities on different MRI sequences, WMH (red arrows) are hyperintense (bright) on T2 FLAIR, but are not hyperintense on T1-weighted images and tend to appear as GM (gray) or CSF (blue) intensity on T1. Thus, T1-based segmentations tend to inflate the GM and CSF volumes in patients with stroke and cerebral small vessel disease. To account for this, ONDRI’s imaging pipeline integrates an additional T2/ FLAIR-based WMH segmentation to correct for this misclassification error (Levy-Cooperman et al., 2008) to produce a normal appearing WM/GM (NAWM/NAGM) volumes.

The NAGM and NAWM volumes can be summed to obtain a measure of parenchymal volume or reported individually for head-size corrected measures to assess potential atrophy. Additionally, a segmentation mask is generated which is used for diffusion tensor imaging (DTI) analyses, where diffusion metrics of the ‘normal appearing’ WM tracts can be separately analysed from the diffusion within the various types of white matter lesions including WMH, lacunar infarcts, and cortical-subcortical stroke lesions. Details of ONDRI DTI analysis pipeline are discussed elsewhere.

The SABRE-LE method segments sCSF and vCSF into separate compartments. The initial T1-based segmentation automatically labels hypointense voxels into a CSF class, and then the ventricles are manually relabelled to a vCSF class by neuroimaging analysts following a standardized procedure. Note that although some vCSF segmentation tools based on standardized templates use smoothing algorithms that reclassify all voxels within the ventricular compartment as ventricles, the SABRELE method does not. With the SABRE-LE method, choroid plexus are not arbitrarily removed or reclassified as CSF and thus remain as part of the overall tissue segmentation. Ventricular volumes are often used as a simple indicator of overall brain atrophy, and have the potential for use as a differential indicator of disease and dementia severity (**Fig.6**) (Nestor et al., 2008; Sapkota et al., 2018; Tavares et al., 2019).

**Fig. 6.**
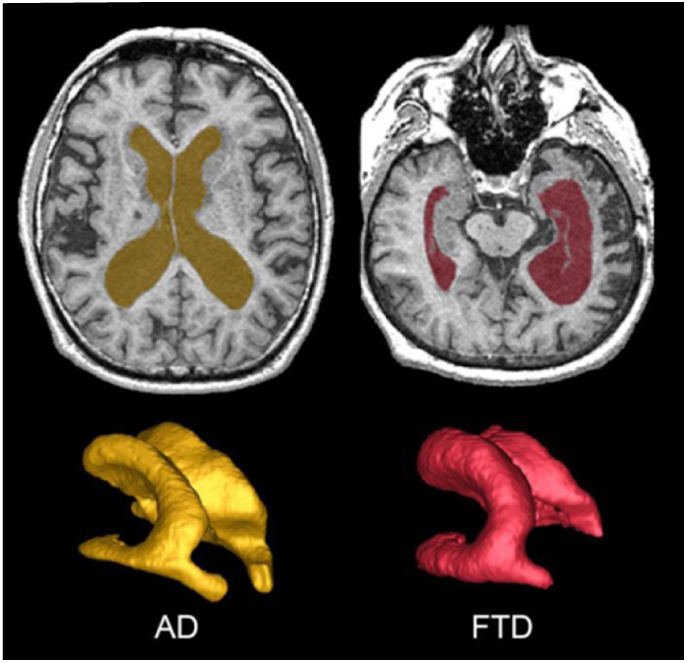
Top row shows axial view of vCSF segmentation overlayed on T1 MRI for patients with AD (left) and FTD (right). Bottom row shows 3D surface volume renderings of the vCSF segmentation. Note the differences in ventricle size and the hemispheric differences between the two neurodegenerative diseases.

### 2.7. White Matter Hyperintensities of presumed vascular origin (WMH)

Also referred to as leukoaraiosis, white matter lesions, subcortical hyperintensities, and even, unidentified bright objects, WMH are radiological anomalies commonly associated with cerebral small vessel disease. Recently, the STandards for ReportIng Vascular changes on nEuroimaging (STRIVE) (Wardlaw et al., 2013) have established a set of criteria that recommends the use of the term *white matter hyperintensities of presumed vascular origin* (WMH), as the standard terminology to refer to these regions of hyperintense (bright) signal found on particular MRI. It is important to note that as previously mentioned, WMH do not appear hyperintense on all types of MRIs and often appear isointense to GM on T1 (**Fig.7**). Additionally, despite the naming convention, it is important to note that WMH are not limited to the white matter regions of the brain, as they are also commonly observed in subcortical GM structures such as the basal ganglia and thalamus. However, to avoid confusion between studies ONDRI recommends the use of the more popular term ‘white matter hyperintensites’.

**Fig. 7.**
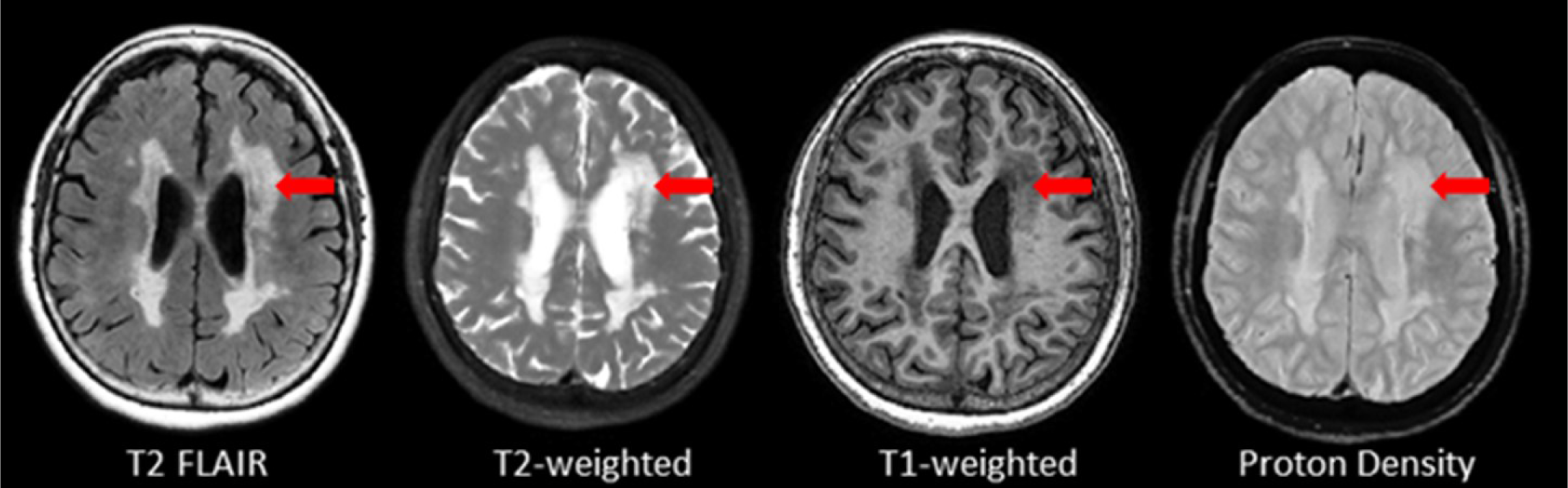
Axial view of various coregistered structural MRI sequences showing the relative intensity differences of WMH. Note that white matter hyperintensities are not hyperintense (i.e. bright) on T1-weighted MRI.

#### 2.7.1. Periventricular (pWMH) and deep white (dWMH) hyper-intensities

Although WMH can be subdivided using SABRE ROIs, the most common regional delineation of WMH is the separation between periventricular (pWMH) and deep white (dWMH). Historically controversial (Barkhof and Scheltens, 2006; Sachdev and Wen, 2005), this concept is based on several theories and research findings which suggest that WMH in close proximity to the ventricles (hence the term ‘peri-ventricular’) have a different pathological etiology (Gouw et al., 2011; Simpson et al., 2007) and are differentially correlated with cognitive/behavior deficits in comparison to the more distal dWMH (despite the confusing fact that pWMH are technically found in deeper white matter than dWMH). Additionally, recent imaging-pathology correlations suggest that a common substrate of pWMH relates to vasogenic edema due to leakage and increased vascular resistance caused by venous collagenosis, a small vessel venular disease of the deep medullary venules (as opposed to the arterial side of the cerebral vasculature) (Black et al., 2009; Keith et al., 2017; Moody et al., 1995). It is also interesting to note that there is no standard consensus in the literature on how to define pWMH vs. dWMH, with some papers using a proportional distance to the dura mater (Decarli et al., 2005a), some using an arbitrary cut-off (typically 13mm from the ventricles) (Sachdev et al., 2008), and others using a 3D connectivity algorithm (Ramirez et al., 2011; van den Heuvel et al., 2006) – the method that is currently supported by ONDRI (see **Fig. 8**).

**Fig. 8.**
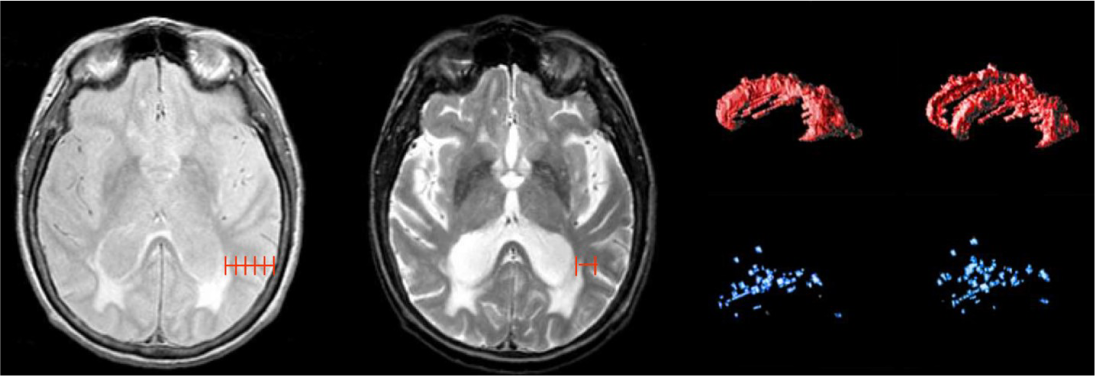
Shows different methods for segmenting periventricular and deep WMH. Left image shows a proportional distance from the ventricular lining to the dura mater; middle image shows an arbitrary distance of 13mm from ventricles, right image shows 3D connectivity algorithm supported by ONDRI, displayed as 3D volume renderings of pWMH (red) and dWMH (blue) shown in sagittal and slightly tilted anterior views.

### 2.8. Lacunes

Lacunes of presumed vascular origin are cystic fluid-filled cavities in the subcortical brain regions (Fisher, 1998; Roman, 2002). They appear hypointense (dark) on T1, hyperintense (bright) on PD and T2, and can appear as a lesion with a hypointense central core surrounded by a hyperintense rim/halo on FLAIR MRI (**Fig. 9, bottom row**). The recent STRIVE criteria (Wardlaw et al., 2013) provides some consensus-based guidelines regarding their definition, however, previous studies have used various terms (eg. ‘*white matter lesions*’, ‘*lacunar infarcts*’, ‘*covert strokes*’) and radiological descriptions to classify these lesions (Potter et al., 2011). Often difficult to differentiate from MRI-visible perivascular spaces (PVS) (next section), lacunes tend to be larger and less linear than PVS. They are associated with increased risk of stroke, dementia, and gait disturbances (Vermeer et al., 2007). It is important to note that due to the poor sensitivity of FLAIR in thalamic regions (Bastos Leite et al., 2004) (**Fig.9, top row**), the ONDRI imaging pipeline integrates an additional T2-based segmentation in order to capture any potential lesions in this subcortical region that may not appear on FLAIR.

**Fig. 9.**
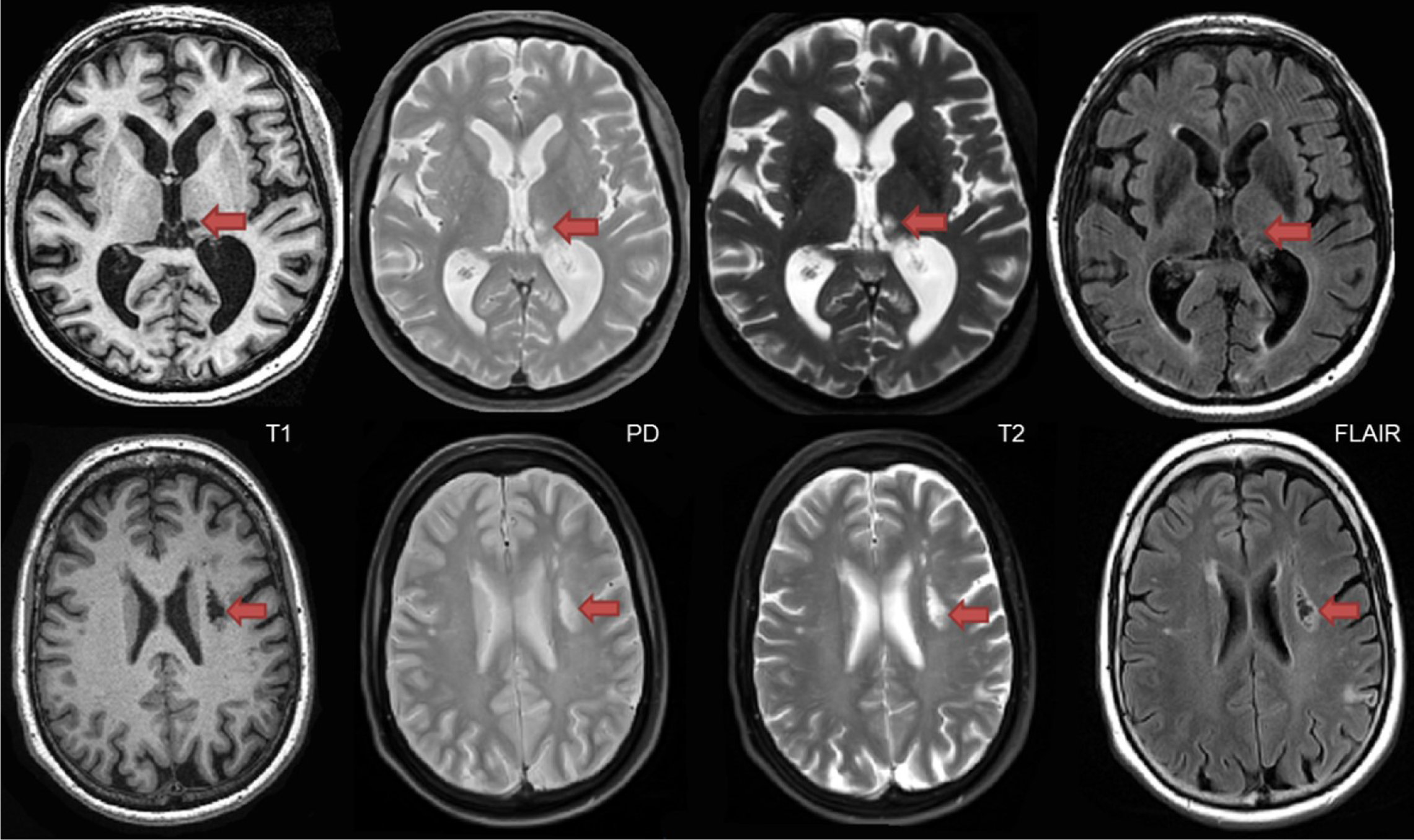
Top row shows a thalamic lacune as it appears on different coregistered MRI, hypointense (dark) on T1, hyperintense on PD-T2, and difficult to detect on FLAIR. In contrast, the bottom row shows a subcortical lacunar infarct that presents with the classic central CSF-like hypointensity with a surrounding hyperintense halo/rim on FLAIR.

### 2.9. MRI-visible (enlarged) Perivascular Spaces (PVS)

Recent studies suggest that the brain utilizes the glymphatic system (Jessen et al., 2015; Rasmussen et al., 2018) to clear fluid and metabolic waste, using a complex series of perivascular channels surrounding the brain’s veins and arteries. It has been suggested that when the perivascular channels are compromised due to aging, disease, or trauma, the perivascular space becomes enlarged and consequently, visible on structural MRI (Ballerini et al., 2018; Ramirez et al., 2016; Tarasoff-Conway et al., 2015). MRI-visible (enlarged) perivascular spaces (PVS) on T2 appear as small (< 3mm diameter), linear, hyperintensities following the course of the vasculature (**Fig. 10**). Additionally, PVS appear hypointense (dark) on T1, isointense to GM on PD (vs. lacunes which are bright on PD), and are very difficult to visualize on 2D FLAIR, particularly in the basal ganglia region. Current research suggests that PVS found in the white matter regions may indicate Cerebral Amyloid Angiopathy (CAA), while PVS in the basal ganglia may be more indicative of hypertensive arteriopathy (Banerjee et al., 2017; A. Charidimou et al., 2015; Charidimou et al., 2014; Martinez-Ramirez et al., 2013). Moreover, recent basic science research and limited clinical evidence supports the theory that clearance of amyloid and other metabolites occurs primarily during deep sleep (Berezuk et al., 2015; Xie et al., 2013).

**Fig. 10.**
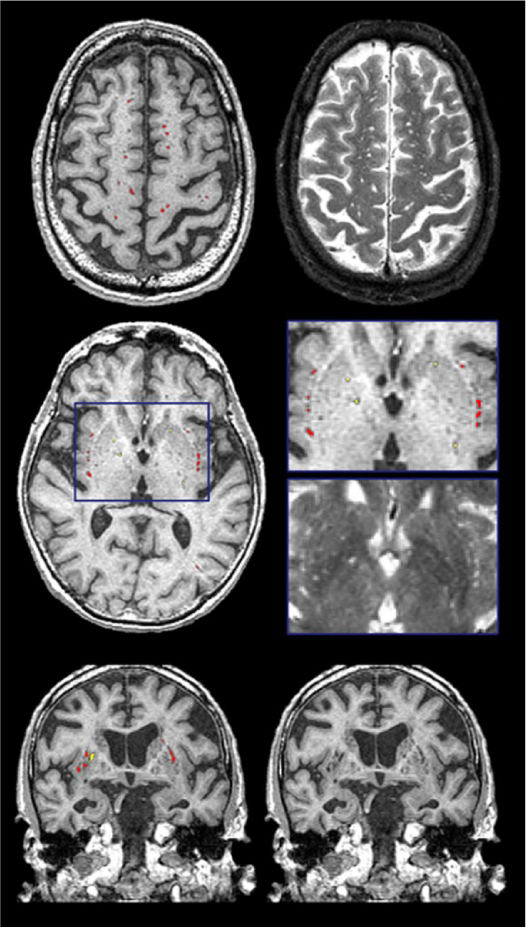
Examples of the PVS segmentation (red & yellow) over-layed onto structural MRI in axial (top 2 rows) and coronal views (bottom row).

Previously referred to as dilated Virchow-Robin spaces, measurement of PVS burden is typically accomplished using visual rating scales under this old naming convention (Adams et al., 2013; Patankar et al., 2005). However, the novel quantitative method supported by ONDRI provides a volumetric measure of PVS. This method has been previously validated with common PVS visual scales and has been used to study AD, normal elderly, and stroke and cerebrovascular disease patients being assessed with sleep polysomnography (Berezuk et al., 2015; Ramirez et al., 2015). Although both lacunes and PVS volumes are segmented automatically using the SABRE-LE pipeline, false positive minimization procedures are manually performed to remove incorrect segmentations and to reallocate PVS to lacunes or vice versa depending on strict intensity and shape-based criteria. Only highly trained neuroimaging analysts achieving ICCs and DICE Similarity Indices (SI) > 0.90 are allowed to perform this procedure. Moreover, a research neuroradiologist (F.G.) was consulted when faced with complex radiological anomalies that were commonly observed in the CVD patient cohort.

### 2.10. Cerebral Microbleeds

Although the SABRE-LE structural pipeline method used by ONDRI does not support a cerebral microbleed (CMB) segmentation algorithm, this brief section has been included to describe this important measure of cerebral small vessel disease burden. In ONDRI, CMB and superficial siderosis burden are being assessed visually by a highly qualified neuroradiologist (S.S.). Cerebral microbleeds (CMB) have been shown to reflect perivascular leakage of red blood cells that can be visualized as low signal intensities (hypointense/dark spots) on T2*-weighted gradient-recalled echo (GRE) (**Fig. 11**) and susceptibility weighted imaging (SWI) (Greenberg et al., 2009). There are two commonly used methods of assessing CMB burden, the Microbleed Anatomical Rating Scale (MARS) (Gregoire et al., 2009) and the Brain Observer MicroBleed Scale (BOMBS) (Cordonnier et al., 2009) visual rating scales. Previous studies have shown that CMB are associated with an increased risk of stroke, intracerebral hemorrhage, cognitive decline, and dementia (Akoudad et al., 2016, 2015; Boyle et al., 2015; Goos et al., 2009; Poels et al., 2010). Differences in anatomical distribution suggest that CMB found in deep centrencephalic brain regions (basal ganglia, thalamus, brain stem) are more closely related to hypertensive arteriopathy (Andreas Charidimou et al., 2015), while lobar CMB are more closely associated with CAA and AD pathology (Cordonnier and van der Flier, 2011; Martinez-Ramirez et al., 2015; Mesker et al., 2011; Pettersen et al., 2008), leading to the development of the Boston criteria for the diagnosis of possible/probable CAA (Boulouis et al., 2016; Smith and Greenberg, 2003).

**Fig. 11.**
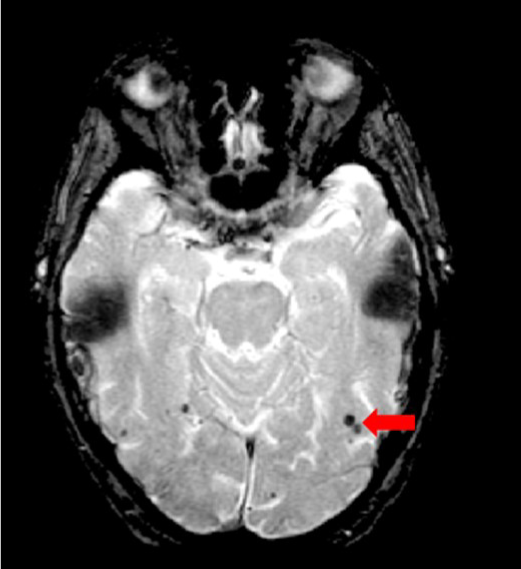
Axial view of iron-sensitive T2* gradient echo (GRE) with red arrow pointing to cerebral microbleeds visualized as hypointensities (dark).

### 2.11. Chronic Stroke

According to recent estimates, stroke is the 2nd most common cause of death worldwide (Feigin et al., 2017) and the second leading cause of dementia (Gorelick et al., 2011). In a 2013 global report, there were approximately 25.7 million stroke survivors, and 7.5 million deaths from ischemic and hemorrhagic stroke (Feigin et al., 2015). In Canada, approximately 62,000 people are treated for stroke and transient ischemic attack. In a series of publications, the Heart and Stroke Foundation Canadian Best Practice Committees have been developing various evidence-based recommendations to address issues regarding: telestroke technologies (Blacquiere et al., 2017); managing transitions of care following stroke (Cameron et al., 2016); mood, cognition, and fatigue following stroke (Lanctôt et al., 2019); hyperacute stroke care (Casaubon et al., 2016); secondary prevention of stroke (Wein et al., 2018); and during pregnancy (Ladhani et al., 2018; Swartz et al., 2018).

Although the term ‘stroke’ may encompass a wide range of clinical criteria (Sacco et al., 2013), the Vascular Cognitive roimaging analysts manually delineate the stroke under the direct supervision of a highly experienced research neuroradiologist (F.G.). This manual delineation is strictly limited to cortical strokes appearing as hyperintense (bright) on FLAIR and hypointense (dark) on T1, although the entire stroke volume often extended into the subcortical regions of the brain (**Fig.13**). Although this total volume does not separate the hypointense necrotic stroke core from the surrounding partially infarcted hyperintense region indicating varying degrees of gliosis and encephalomalacia, future automatic segmentation techniques are currently being tested in ONDRI to include this sub-segmentation.

**Fig. 12.**
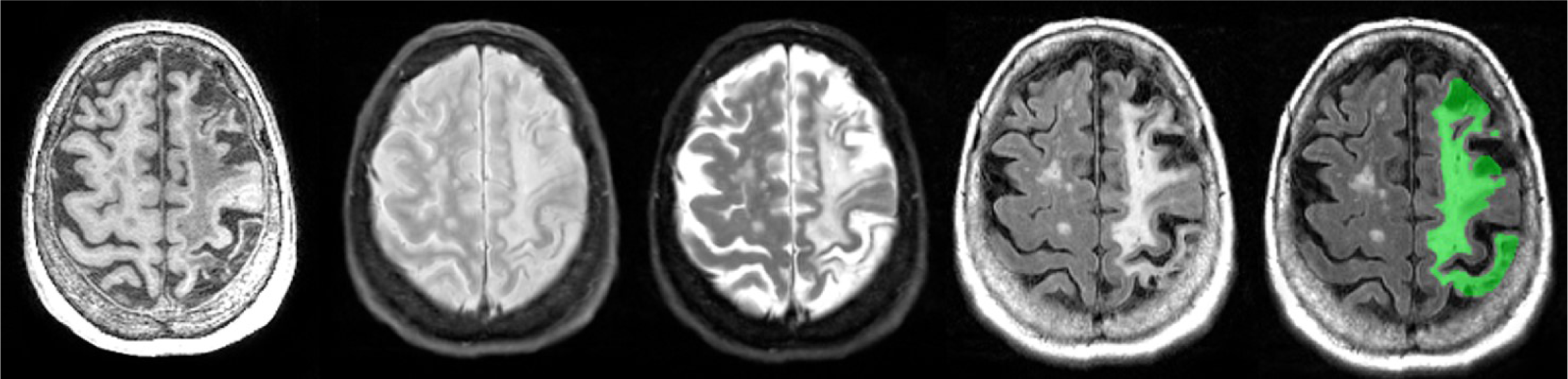
Axial view of coregistered structural MRI sequences (left to right): T1, PD, T2, FLAIR. Images illustrate relative intensity differences of a large cortico-subcortical stroke lesion across various types of MRI. The last pane shows ONDRI’s manual segmentation of the entire stroke core and surrounding hyperintense partially infarcted tissue volume in green.

**Fig. 13.**
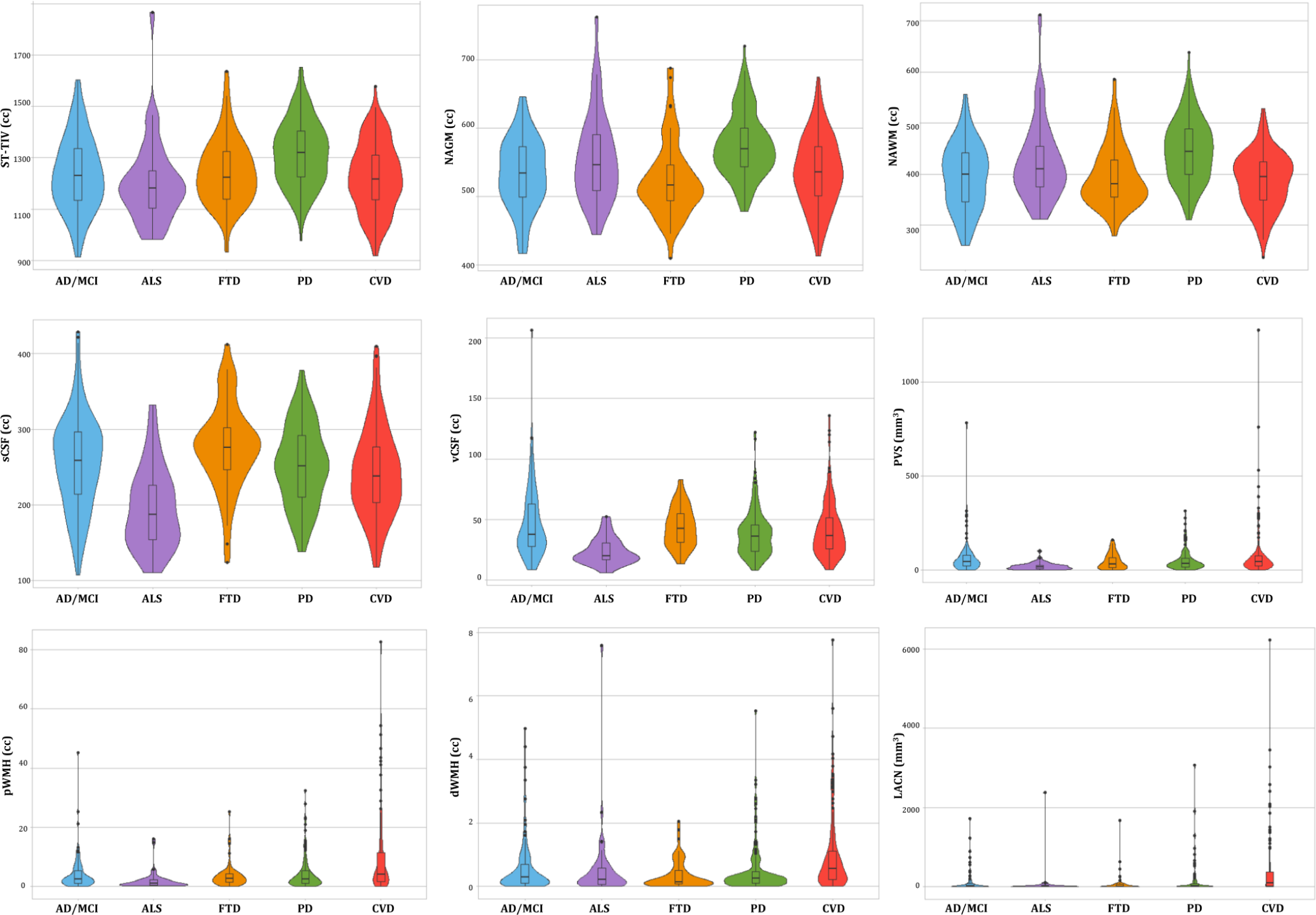
Descriptive violin plots of the ONDRI disease cohorts showing median and interquartile range volumetrics for whole brain supratentorial total intracranial volume (ST-TIV), normal appearing gray matter (NAGM), normal appearing white matter (NAWM), sulcal cerebrospinal fluid (sCSF), ventricular CSF (vCSF), MRI-visible enlarged perivascular space (PVS) volumes, periventricular white matter hyperintensities (pWMH), deep WMH (dWMH), and lacunes (LACN).

## 3. RECOMMENDATIONS FOR REPORTING & ANALYSIS

Here we provide some general guidelines for reporting and analysis that can be useful for researchers wishing to use ONDRI data.

First and foremost, when reporting data for characterization of the sample being analysed, we recommend that the original raw volumes are reported in tables for transparency and between-study comparisons; however, statistical analyses should generally be performed on head-size corrected volumes. Head size correction accounts for individual variations in intracranial capacity and sex-related differences in head size (Decarli et al., 2005b). Additionally, depending on the research question, the volume of interest (i.e., NAWM, NAGM, CSF, or WMH) could also be reported as a proportion of the total volume within each SABRE region, or they can be reported as a proportion of the total head-size (ST-TIV) for age-independent normalization/correction. The version or date of the data release should also be reported.

There are several ways that WMH can be analysed and it depends on the research question in mind. The simplest approach is to sum the dWMH and pWMH, which results in a whole brain measure of small vessel disease burden. Regional anal-Impairment (VCI) (Hachinski et al., 2006) inclusion-exclusion criteria for ONDRI CVD patients was limited to mild-moderate ischemic stroke patients, defined by a Modified Rankin Scale (MRS) (van Swieten et al., 1988) score of 0 to 3. It is important to note that although there are a number of imaging techniques used to measure acute stroke in the early stages (within a couple of hours of stroke), the MRI methods applied to ONDRI CVD patients are measures of post-stroke lesions, often referred to as *chronic stroke*, with structural MRI acquired > 3 months post ischemic stroke event.

As there are currently no reliable automatic ways to quantify the range of cortico-subcortical stroke lesions, ONDRI neu-yses of WMH can also be performed to assess WMH burden within a SABRE ROI. Additionally, WMH within different ROIs can be combined by simply summing the volumes from different SABRE regions to generate a larger ROI (eg. sum all pWMH and dWMH volumes within all frontal SABRE brain region parcellations using the Frontal Lobe Codes shown on Table 1).

It is important to note that many measures of cerebral small vessel disease, such as pWMH and dWMH, are typically non-normally distributed (Carmichael et al., 2010), often intercorrelated (Decarli et al., 2005a), are known to be age-related (Pantoni, 2010), and commonly associated with vascular risk factors such as hypertension (Basile et al., 2006). Thus, careful attention to these factors and previous research findings highlight ONDRI’s recommendation to consider these additional factors when analyzing imaging-based markers of cerebral small vessel disease. Given the skewed, non-normal distribution of WMH (even after head-size correction), WMH volumes are typically transformed (eg. log) prior to standard parametric analyses. For this reason, approaches designed to deal with complex distributions should be considered (Nguyen and Holmes, 2019).

Since the pipeline automatically segments lesions in the periventricular region from the deep white regions, the lacunar volumes are also provided in this manner. While some future studies may argue a pathophysiological difference between these two locations of lacunar presentation, there are currently limited studies to suggest this anatomical delineation. Given this, we recommend that the two volumes be summed together prior to analysis. Interestingly, lacunes and PVS volumes are not typically head-size corrected in the clinical/scientific literature, however, age, sex, WMH, and a measure of brain atrophy (eg. BPF or vCSF), and proper accounting of vascular risk are recommended covariates when analyzing lacunes and PVS (Zhu et al., 2011, 2010). Note that in many publications, lacunes and PVS are reported as counts (i.e. number of), because they are often measured using visual rating methods that require the user to count the number of lacunes or PVS observed on an MRI – often leading to wide variations in definitions and conflicting findings in the literature (Francis et al., 2019; Potter et al., 2011). Since the lacunes and PVS in ON-DRI are quantified using segmentation based imaging analyses, PVS and lacunar volumes are provided rather than counts.

Finally, any analyses using ONDRI’s CVD cohort should consider the common comorbidities of depression, obstructive sleep apnea, and cognitive impairment (Swartz et al., 2016), as well the subcortical silent brain infarcts/lacunes, WMH, and potentially, CMB, which have recently been acknowledged as playing an important role in primary stroke prevention (Smith et al., 2017).

## 4. RESULTS AND CONCLUSION

Of the n=520 patients with MRI acquired, the ONDRI neuroimaging pipeline was unable to process n=1 FTD patient due to extreme motion artifact (despite 2 baseline attempts on 2 separate occasions), and n=6 CVD patients due to poor imaging quality (n=3 FLAIR not usable, n=2 T1 not usable, n=1 PD/T2 not acquired). To illustrate whole brain data extraction volumetric results from this pipeline, neuroimaging summary statistics for each ONDRI disease cohort are summarized on **Table 2**, and descriptive violin plots showing median and interquartile ranges are provided for whole brain ST-TIV, NAGM, NAWM, sCSF, vCSF, pWMH, and dWMH PVS, and LACN are displayed on **Fig.13**. Stroke volumes were not graphed due to the limited number of ONDRI patients with cortico-subcortical stroke lesions.

ONDRI is the first multi-site, multiple assessment platform study examining several neurodegenerative and neurovascular diseases using a harmonized protocol that includes standardized structural neuroimaging. The wide range of complex, and often overlapping, brain pathologies represented in this cohort of neurodegenerative patients included a number of comorbid cerebral small vessel disease markers, cortico-subcortical stroke lesions, combined with focal and global atrophy, posing significant challenges to common imaging analysis tools. In this paper, we presented the neuroimaging pipeline methods implemented in ONDRI that were used to overcome many of these challenges. To further ensure a high level of data quality, the volumetric data generated by the ONDRI structural neuroimaging team were further subjected to comprehensive quality control analysis pipelines including a novel multivariate outlier detection algorithm developed by the ONDRI neuroinformatics group for identification of anomalous observations (Beaton et al., 2019; Sunderland et al., 2019). Future work will include generating longitudinal measures that will also be made publicly available. As the neuroimaging data are combined with releases from ONDRI’s clinical, neuropsychology, genomics, eye tracking, gait and balance, ocular, and neuropathology platforms, it becomes evident that ONDRI is a gold mine of data opening the door to an unprecedented broad range of crossplatform analyses resulting in numerous opportunities for discovery and advances in diagnosis, prognosis, outcomes, and care of neurodegenerative diseases.

## ACKNOWLEDGEMENTS

We would like to thank the ONDRI participants for the time, consent, and participation in our study. Thank you to the L.C. Campbell Foundation, and the analysts and software developers in the LC Campbell Cognitive Neurology research team who have contributed to the ONDRI imaging analysis, including Edward Ntiri, Hassan Akhavein, Parisa Mojiri, Kirstin Walker, Rita Meena, Pugaliya Puveendrakumaran, Courtney Berezuk, and Alicia McNeely.

---

**Supplementary Table 1.**
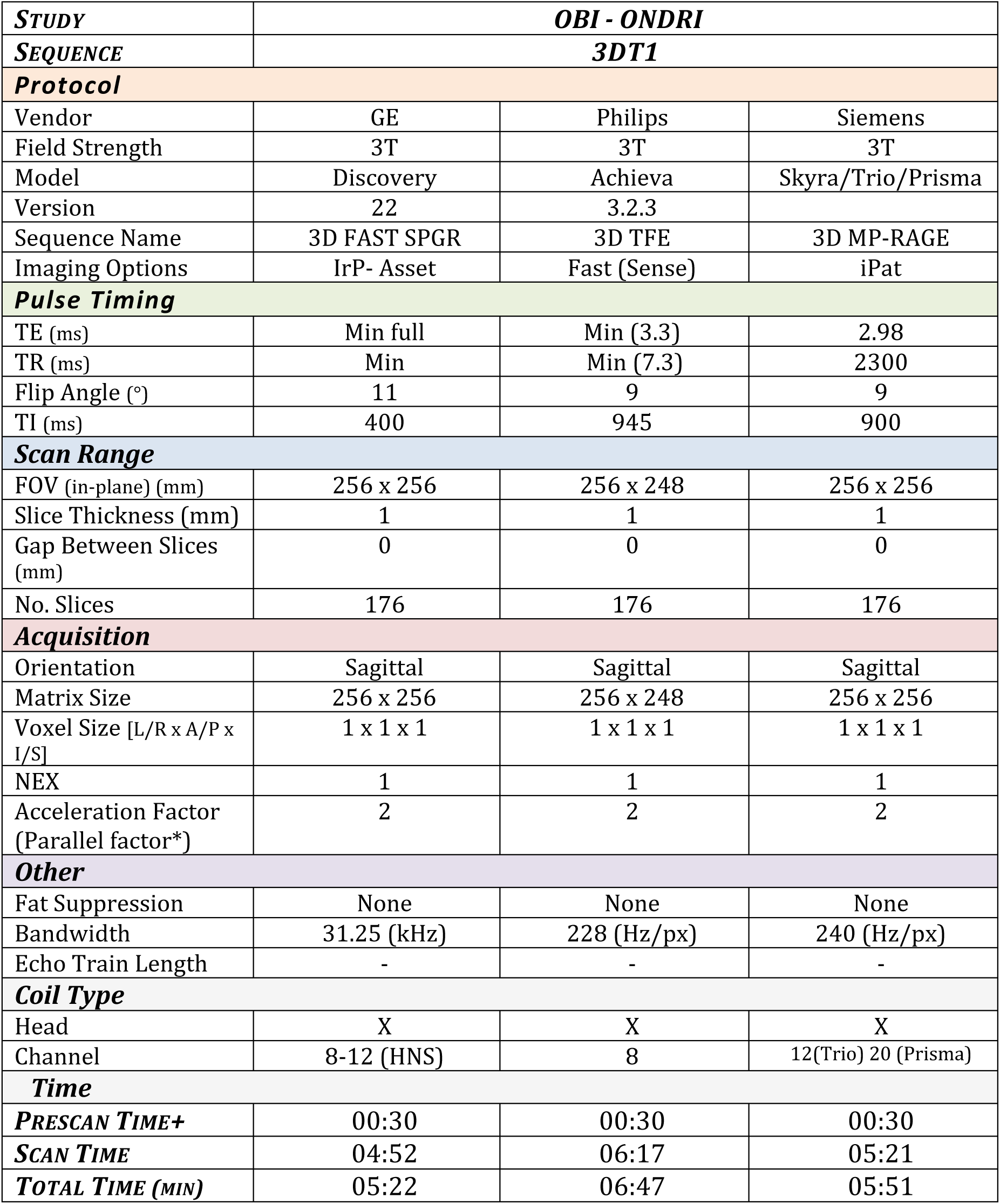
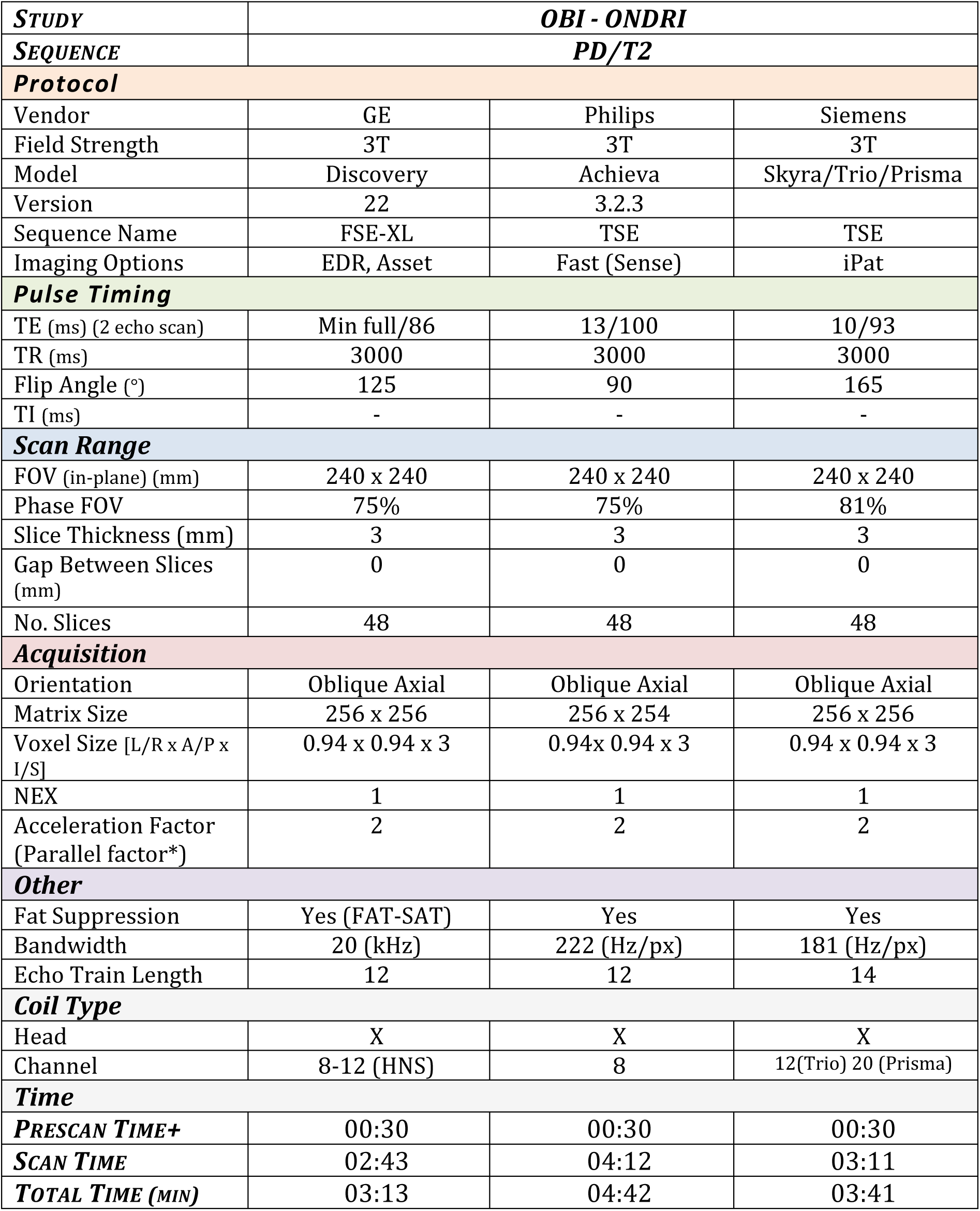
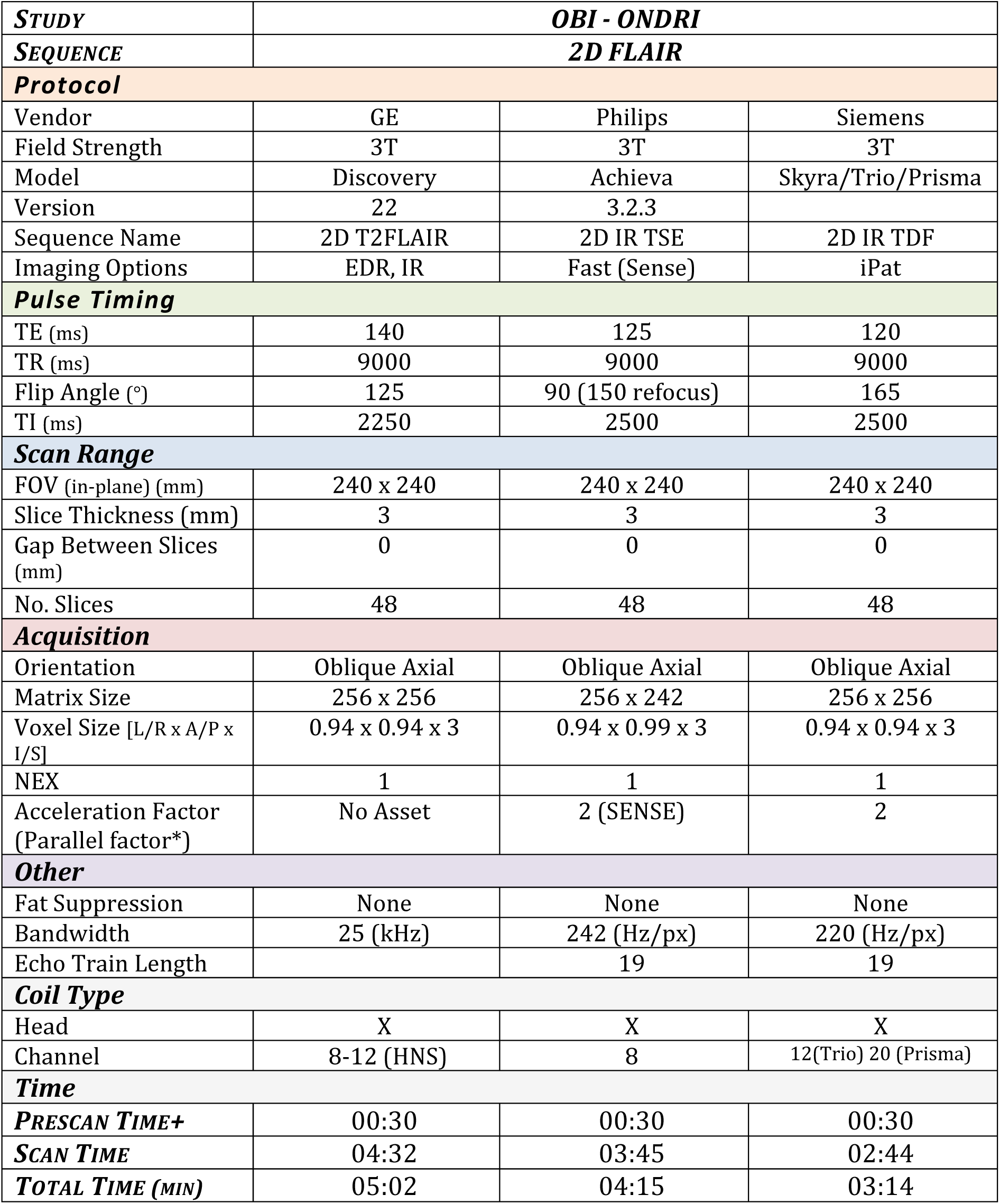
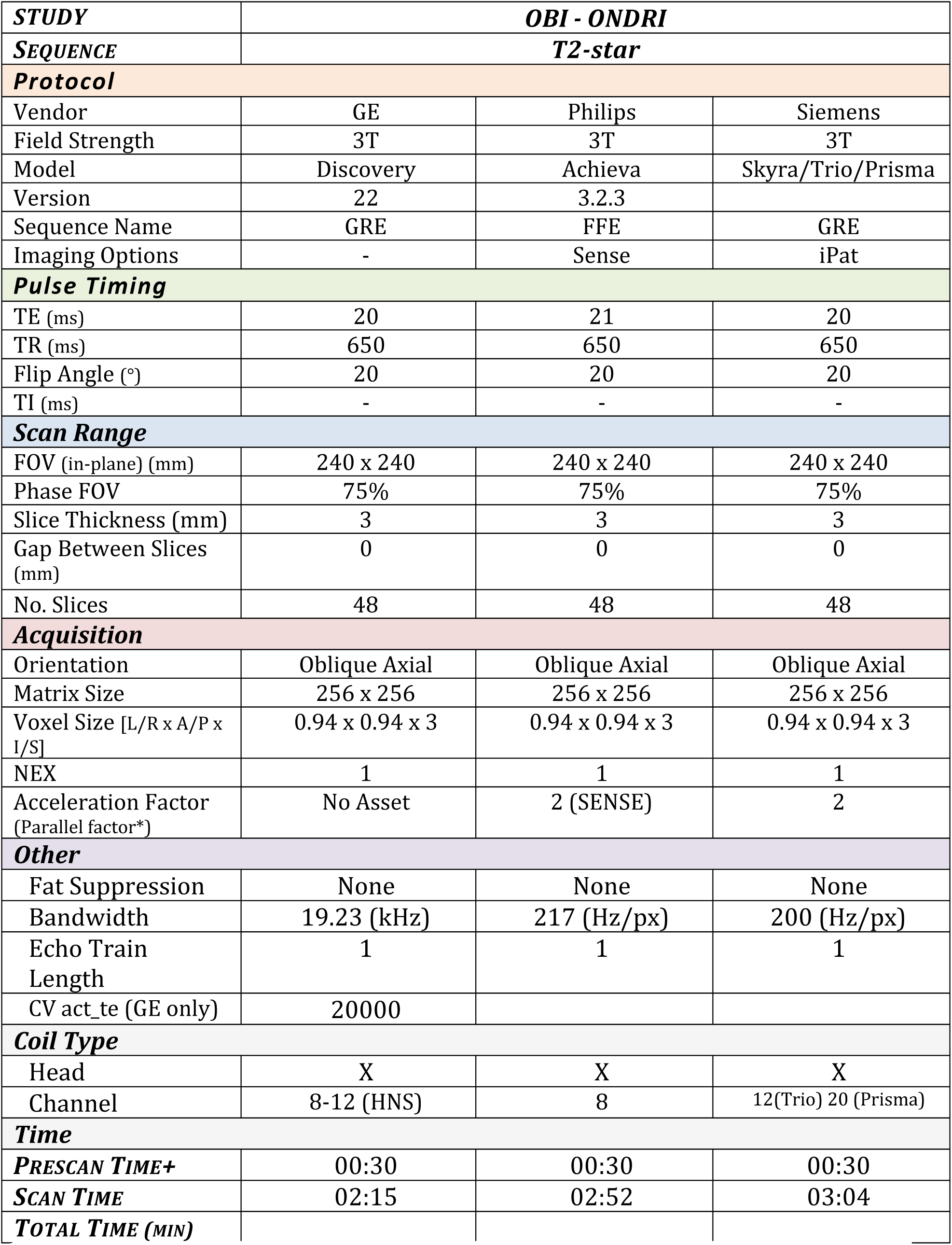
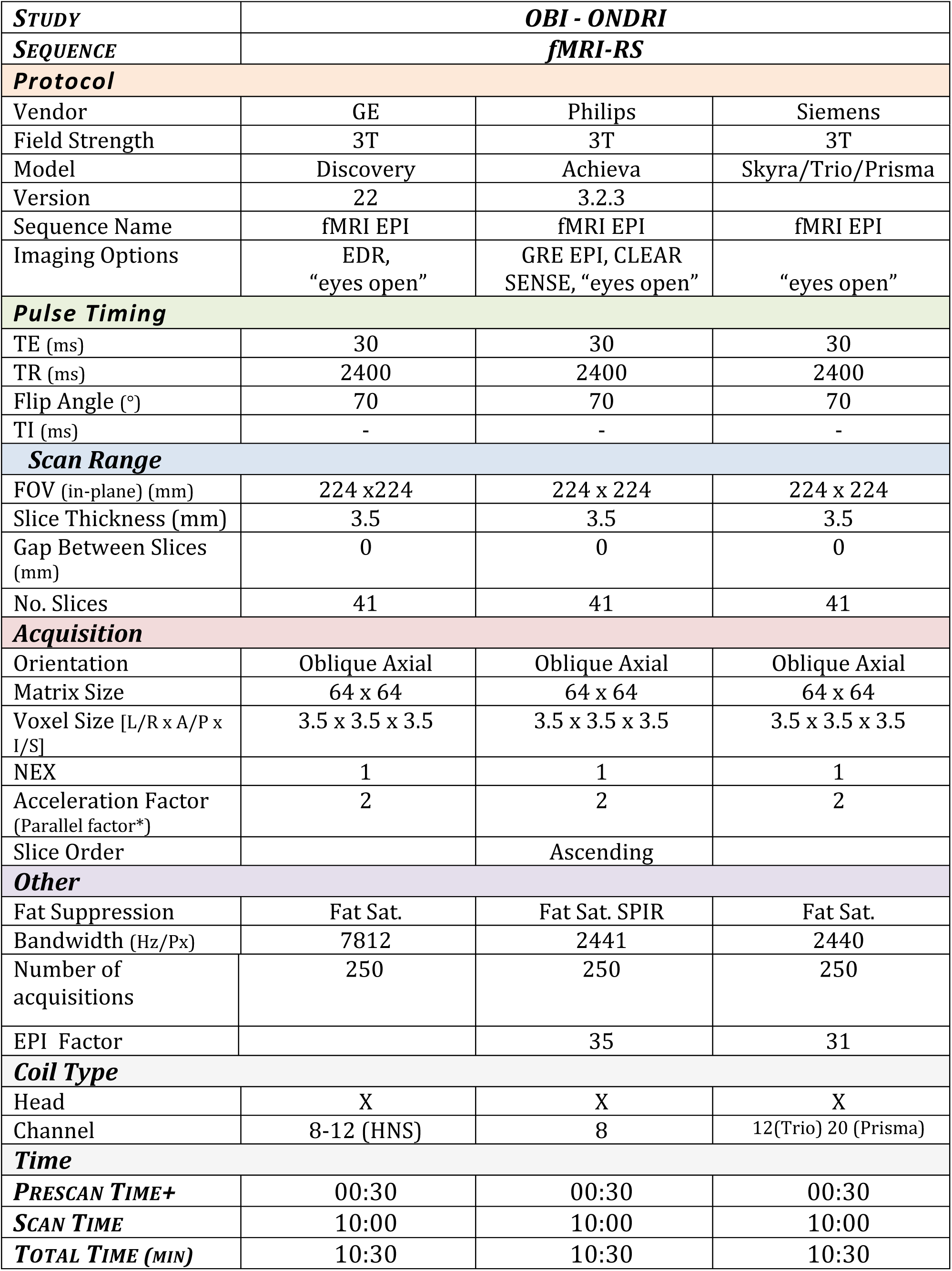
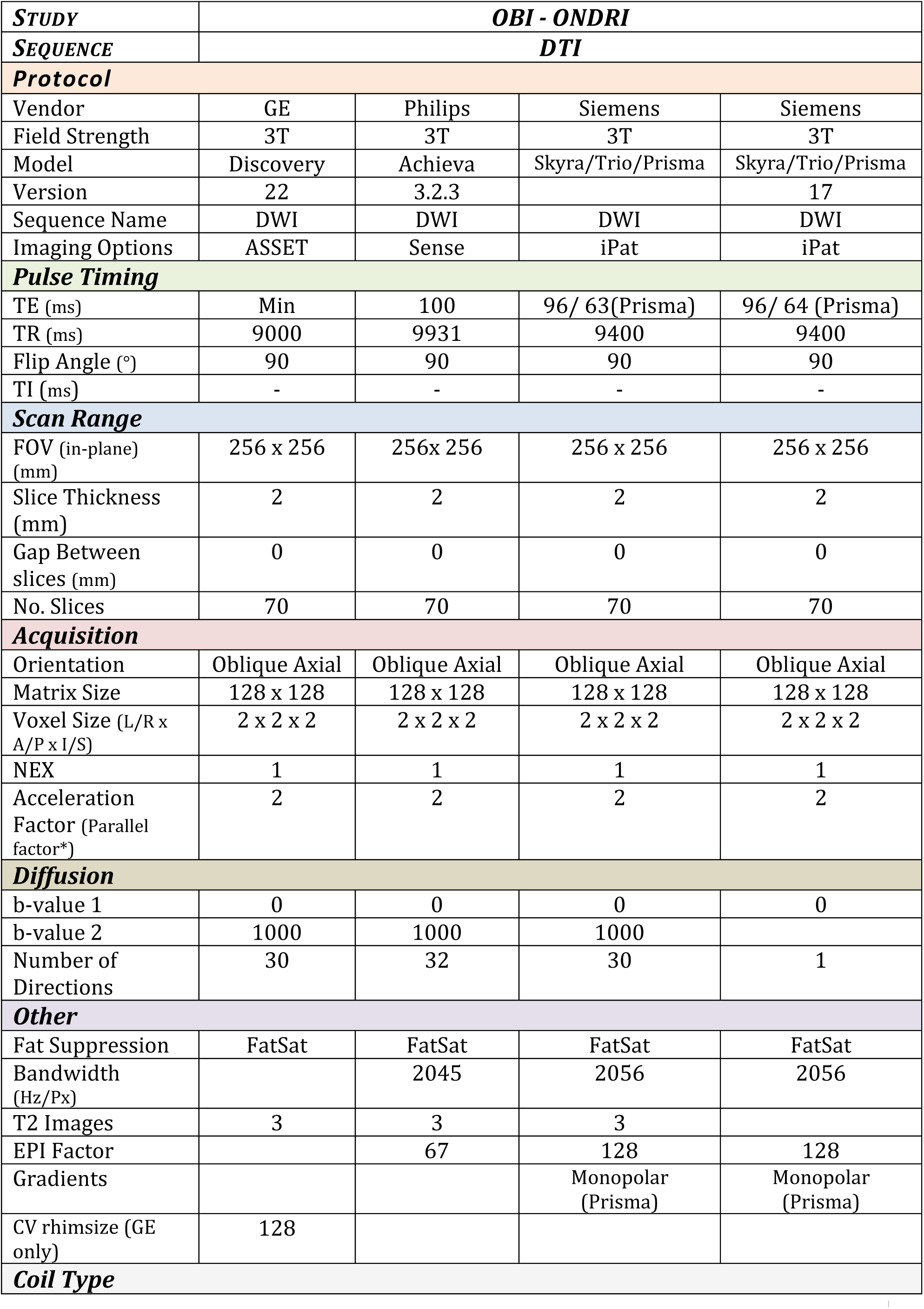
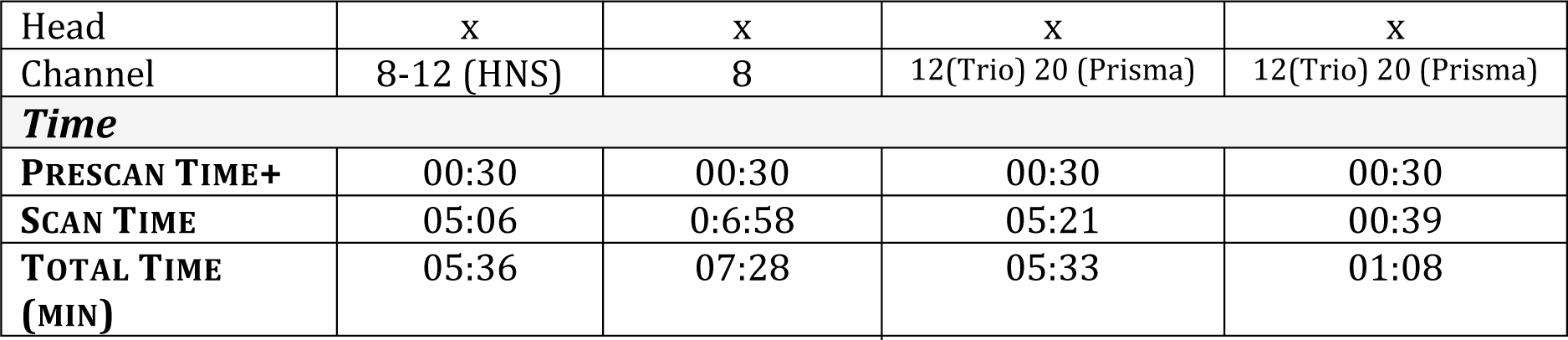
MRI Acquisition Protocols.

|  |  |  |  |
| --- | --- | --- | --- |
| **STUDY** | **OBI - ONDRI** |  |  |
| **SEQUENCE** | **3DT1** |  |  |
| **Protocol** |  |  |  |
| Vendor | GE | Philips | Siemens |
| Field Strength | 3T | 3T | 3T |
| Model | Discovery | Achieva | Skyra/Trio/Prisma |
| Version | 22 | 3.2.3 |  |
| Sequence Name | 3D FAST SPGR | 3D TFE | 3D MP-RAGE |
| Imaging Options | IrP- Asset | Fast (Sense) | iPat |
| **Pulse Timing** |  |  |  |
| TE (ms) | Min full | Min (3.3) | 2.98 |
| TR (ms) | Min | Min (7.3) | 2300 |
| Flip Angle (°) | 11 | 9 | 9 |
| TI (ms) | 400 | 945 | 900 |
| **Scan Range** |  |  |  |
| FOV (in-plane) (mm) | 256 x 256 | 256 x 248 | 256 x 256 |
| Slice Thickness (mm) | 1 | 1 | 1 |
| Gap Between Slices (mm) | 0 | 0 | 0 |
| No. Slices | 176 | 176 | 176 |
| **Acquisition** |  |  |  |
| Orientation | Sagittal | Sagittal | Sagittal |
| Matrix Size | 256 x 256 | 256 x 248 | 256 x 256 |
| Voxel Size [L/R x A/P x I/S] | 1 x 1 x 1 | 1 x 1 x 1 | 1 x 1 x 1 |
| NEX | 1 | 1 | 1 |
| Acceleration Factor (Parallel factor*) | 2 | 2 | 2 |
| **Other** |  |  |  |
| Fat Suppression | None | None | None |
| Bandwidth | 31.25 (kHz) | 228 (Hz/px) | 240 (Hz/px) |
| Echo Train Length | - | - | - |
| **Coil Type** |  |  |  |
| Head | X | X | X |
| Channel | 8-12 (HNS) | 8 | 12(Trio) 20 (Prisma) |
| **Time** |  |  |  |
| **PRESCAN TIME+** | 00:30 | 00:30 | 00:30 |
| **SCAN TIME** | 04:52 | 06:17 | 05:21 |
| **TOTAL TIME (MIN)** | 05:22 | 06:47 | 05:51 |
| **STUDY** | **OBI - ONDRI** |  |  |
| **SEQUENCE** | **PD/T2** |  |  |
| **Protocol** |  |  |  |
| Vendor | GE | Philips | Siemens |
| Field Strength | 3T | 3T | 3T |
| Model | Discovery | Achieva | Skyra/Trio/Prisma |
| Version | 22 | 3.2.3 |  |
| Sequence Name | FSE-XL | TSE | TSE |
| Imaging Options | EDR, Asset | Fast (Sense) | iPat |
| **Pulse Timing** |  |  |  |
| TE (ms) (2 echo scan) | Min full/86 | 13/100 | 10/93 |
| TR (ms) | 3000 | 3000 | 3000 |
| Flip Angle (°) | 125 | 90 | 165 |
| TI (ms) | - | - | - |
| **Scan Range** |  |  |  |
| FOV (in-plane) (mm) | 240 x 240 | 240 x 240 | 240 x 240 |
| Phase FOV | 75% | 75% | 81% |
| Slice Thickness (mm) | 3 | 3 | 3 |
| Gap Between Slices (mm) | 0 | 0 | 0 |
| No. Slices | 48 | 48 | 48 |
| **Acquisition** |  |  |  |
| Orientation | Oblique Axial | Oblique Axial | Oblique Axial |
| Matrix Size | 256 x 256 | 256 x 254 | 256 x 256 |
| Voxel Size [L/R x A/P x I/S] | 0.94 x 0.94 x 3 | 0.94x 0.94 x 3 | 0.94 x 0.94 x 3 |
| NEX | 1 | 1 | 1 |
| Acceleration Factor (Parallel factor*) | 2 | 2 | 2 |
| **Other** |  |  |  |
| Fat Suppression | Yes (FAT-SAT) | Yes | Yes |
| Bandwidth | 20 (kHz) | 222 (Hz/px) | 181 (Hz/px) |
| Echo Train Length | 12 | 12 | 14 |
| **Coil Type** |  |  |  |
| Head | X | X | X |
| Channel | 8-12 (HNS) | 8 | 12(Trio) 20 (Prisma) |
| **Time** |  |  |  |
| **PRESCAN TIME+** | 00:30 | 00:30 | 00:30 |
| **SCAN TIME** | 02:43 | 04:12 | 03:11 |
| **TOTAL TIME (MIN)** | 03:13 | 04:42 | 03:41 |
| **STUDY** | **OBI - ONDRI** |  |  |
| **SEQUENCE** | **2D FLAIR** |  |  |
| **Protocol** |  |  |  |
| Vendor | GE | Philips | Siemens |
| Field Strength | 3T | 3T | 3T |
| Model | Discovery | Achieva | Skyra/Trio/Prisma |
| Version | 22 | 3.2.3 |  |
| Sequence Name | 2D T2FLAIR | 2D IR TSE | 2D IR TDF |
| Imaging Options | EDR, IR | Fast (Sense) | iPat |
| **Pulse Timing** |  |  |  |
| TE (ms) | 140 | 125 | 120 |
| TR (ms) | 9000 | 9000 | 9000 |
| Flip Angle (°) | 125 | 90 (150 refocus) | 165 |
| TI (ms) | 2250 | 2500 | 2500 |
| **Scan Range** |  |  |  |
| FOV (in-plane) (mm) | 240 x 240 | 240 x 240 | 240 x 240 |
| Slice Thickness (mm) | 3 | 3 | 3 |
| Gap Between Slices (mm) | 0 | 0 | 0 |
| No. Slices | 48 | 48 | 48 |
| **Acquisition** |  |  |  |
| Orientation | Oblique Axial | Oblique Axial | Oblique Axial |
| Matrix Size | 256 x 256 | 256 x 242 | 256 x 256 |
| Voxel Size [L/R x A/P x I/S] | 0.94 x 0.94 x 3 | 0.94 x 0.99 x 3 | 0.94 x 0.94 x 3 |
| NEX | 1 | 1 | 1 |
| Acceleration Factor (Parallel factor*) | No Asset | 2 (SENSE) | 2 |
| **Other** |  |  |  |
| Fat Suppression | None | None | None |
| Bandwidth | 25 (kHz) | 242 (Hz/px) | 220 (Hz/px) |
| Echo Train Length |  | 19 | 19 |
| **Coil Type** |  |  |  |
| Head | X | X | X |
| Channel | 8-12 (HNS) | 8 | 12(Trio) 20 (Prisma) |
| **Time** |  |  |  |
| **PRESCAN TIME+** | 00:30 | 00:30 | 00:30 |
| **SCAN TIME** | 04:32 | 03:45 | 02:44 |
| **TOTAL TIME (MIN)** | 05:02 | 04:15 | 03:14 |
| **STUDY** | **OBI - ONDRI** |  |  |
| **SEQUENCE** | **T2-star** |  |  |
| **Protocol** |  |  |  |
| Vendor | GE | Philips | Siemens |
| Field Strength | 3T | 3T | 3T |
| Model | Discovery | Achieva | Skyra/Trio/Prisma |
| Version | 22 | 3.2.3 |  |
| Sequence Name | GRE | FFE | GRE |
| Imaging Options | - | Sense | iPat |
| **Pulse Timing** |  |  |  |
| TE (ms) | 20 | 21 | 20 |
| TR (ms) | 650 | 650 | 650 |
| Flip Angle (°) | 20 | 20 | 20 |
| TI (ms) | - | - | - |
| **Scan Range** |  |  |  |
| FOV (in-plane) (mm) | 240 x 240 | 240 x 240 | 240 x 240 |
| Phase FOV | 75% | 75% | 75% |
| Slice Thickness (mm) | 3 | 3 | 3 |
| Gap Between Slices (mm) | 0 | 0 | 0 |
| No. Slices | 48 | 48 | 48 |
| **Acquisition** |  |  |  |
| Orientation | Oblique Axial | Oblique Axial | Oblique Axial |
| Matrix Size | 256 x 256 | 256 x 256 | 256 x 256 |
| Voxel Size [L/R x A/P x I/S] | 0.94 x 0.94 x 3 | 0.94 x 0.94 x 3 | 0.94 x 0.94 x 3 |
| NEX | 1 | 1 | 1 |
| Acceleration Factor (Parallel factor*) | No Asset | 2 (SENSE) | 2 |
| **Other** |  |  |  |
| Fat Suppression | None | None | None |
| Bandwidth | 19.23 (kHz) | 217 (Hz/px) | 200 (Hz/px) |
| Echo Train Length | 1 | 1 | 1 |
| CV act_te (GE only) | 20000 |  |  |
| **Coil Type** |  |  |  |
| Head | X | X | X |
| Channel | 8-12 (HNS) | 8 | 12(Trio) 20 (Prisma) |
| **Time** |  |  |  |
| **PRESCAN TIME+** | 00:30 | 00:30 | 00:30 |
| **SCAN TIME** | 02:15 | 02:52 | 03:04 |
| **TOTAL TIME (MIN)** |  |  |  |
| **STUDY** | **OBI - ONDRI** |  |  |
| **SEQUENCE** | **fMRI-RS** |  |  |
| **Protocol** |  |  |  |
| Vendor | GE | Philips | Siemens |
| Field Strength | 3T | 3T | 3T |
| Model | Discovery | Achieva | Skyra/Trio/Prisma |
| Version | 22 | 3.2.3 |  |
| Sequence Name | fMRI EPI | fMRI EPI | fMRI EPI |
| Imaging Options | EDR,<br>“eyes open” | GRE EPI, CLEAR<br>SENSE, “eyes open” | “eyes open” |
| **Pulse Timing** |  |  |  |
| TE (ms) | 30 | 30 | 30 |
| TR (ms) | 2400 | 2400 | 2400 |
| Flip Angle (°) | 70 | 70 | 70 |
| TI (ms) | - | - | - |
| **Scan Range** |  |  |  |
| FOV (in-plane) (mm) | 224 x224 | 224 x 224 | 224 x 224 |
| Slice Thickness (mm) | 3.5 | 3.5 | 3.5 |
| Gap Between Slices (mm) | 0 | 0 | 0 |
| No. Slices | 41 | 41 | 41 |
| **Acquisition** |  |  |  |
| Orientation | Oblique Axial | Oblique Axial | Oblique Axial |
| Matrix Size | 64 x 64 | 64 x 64 | 64 x 64 |
| Voxel Size [L/R x A/P x I/S] | 3.5 x 3.5 x 3.5 | 3.5 x 3.5 x 3.5 | 3.5 x 3.5 x 3.5 |
| NEX | 1 | 1 | 1 |
| Acceleration Factor (Parallel factor*) | 2 | 2 | 2 |
| Slice Order |  | Ascending |  |
| **Other** |  |  |  |
| Fat Suppression | Fat Sat. | Fat Sat. SPIR | Fat Sat. |
| Bandwidth (Hz/Px) | 7812 | 2441 | 2440 |
| Number of acquisitions | 250 | 250 | 250 |
| EPI Factor |  | 35 | 31 |
| **Coil Type** |  |  |  |
| Head | X | X | X |
| Channel | 8-12 (HNS) | 8 | 12(Trio) 20 (Prisma) |
| **Time** |  |  |  |
| **PRESCAN TIME+** | 00:30 | 00:30 | 00:30 |
| **SCAN TIME** | 10:00 | 10:00 | 10:00 |
| **TOTAL TIME (MIN)** | 10:30 | 10:30 | 10:30 |

|  |  |  |  |  |
| --- | --- | --- | --- | --- |
| **STUDY** | **OBI - ONDRI** |  |  |  |
| **SEQUENCE** | **DTI** |  |  |  |
| **Protocol** |  |  |  |  |
| Vendor | GE | Philips | Siemens | Siemens |
| Field Strength | 3T | 3T | 3T | 3T |
| Model | Discovery | Achieva | Skyra/Trio/Prisma | Skyra/Trio/Prisma |
| Version | 22 | 3.2.3 |  | 17 |
| Sequence Name | DWI | DWI | DWI | DWI |
| Imaging Options | ASSET | Sense | iPat | iPat |
| **Pulse Timing** |  |  |  |  |
| TE (ms) | Min | 100 | 96/ 63(Prisma) | 96/ 64 (Prisma) |
| TR (ms) | 9000 | 9931 | 9400 | 9400 |
| Flip Angle (°) | 90 | 90 | 90 | 90 |
| TI (ms) | - | - | - | - |
| **Scan Range** |  |  |  |  |
| FOV (in-plane) (mm) | 256 x 256 | 256x 256 | 256 x 256 | 256 x 256 |
| Slice Thickness (mm) | 2 | 2 | 2 | 2 |
| Gap Between slices (mm) | 0 | 0 | 0 | 0 |
| No. Slices | 70 | 70 | 70 | 70 |
| **Acquisition** |  |  |  |  |
| Orientation | Oblique Axial | Oblique Axial | Oblique Axial | Oblique Axial |
| Matrix Size | 128 x 128 | 128 x 128 | 128 x 128 | 128 x 128 |
| Voxel Size (L/R x A/P x I/S) | 2 x 2 x 2 | 2 x 2 x 2 | 2 x 2 x 2 | 2 x 2 x 2 |
| NEX | 1 | 1 | 1 | 1 |
| Acceleration Factor (Parallel factor*) | 2 | 2 | 2 | 2 |
| **Diffusion** |  |  |  |  |
| b-value 1 | 0 | 0 | 0 | 0 |
| b-value 2 | 1000 | 1000 | 1000 |  |
| Number of Directions | 30 | 32 | 30 | 1 |
| **Other** |  |  |  |  |
| Fat Suppression | FatSat | FatSat | FatSat | FatSat |
| Bandwidth (Hz/Px) |  | 2045 | 2056 | 2056 |
| T2 Images | 3 | 3 | 3 |  |
| EPI Factor |  | 67 | 128 | 128 |
| Gradients |  |  | Monopolar (Prisma) | Monopolar (Prisma) |
| CV rhimsize (GE only) | 128 |  |  |  |
| **Coil Type** |  |  |  |  |
| Head | x | x | x | x |
| Channel | 8-12 (HNS) | 8 | 12(Trio) 20 (Prisma) | 12(Trio) 20 (Prisma) |
| **Time** |  |  |  |  |
| **PRESCAN TIME+** | 00:30 | 00:30 | 00:30 | 00:30 |
| **SCAN TIME** | 05:06 | 0:6:58 | 05:21 | 00:39 |
| **TOTAL TIME**<br>**(MIN)** | 05:36 | 07:28 | 05:33 | 01:08 |

## Core Protocol Scan Time Estimates

|  | *GE* | *Philips* | *Siemens* |
| --- | --- | --- | --- |
| Set up | 05:00 | 05:00 | 05:00 |
| 3DT1 | 05:22 | 06:47 | 05:51 |
| PD/T2 | 02:49 | 04:42 | 03:41 |
| 2D FLAIR | 05:02 | 04:15 | 03:14 |
| T2-star *or* T2-star_PM *or* T2-star_PMRI | 03:45 | 03:22 | 03:34 |
| RS-fMRI | 10:30 | 10:30 | 10:30 |
| DTI | 06:30 | 07:28 | 06:41 |
| Set down | 02:30 | 02:30 | 02:30 |
| **Total scan time** | 33:58 | 37:04 | 33:31 |
| **Total session time** | **41:28** | **44:34** | **41:01** |

